# Kinetochore-centrosome feedback linking CENP-E and Aurora kinases controls chromosome congression

**DOI:** 10.1101/2024.09.29.614573

**Authors:** Kruno Vukušić, Iva M. Tolić

## Abstract

Chromosome congression is crucial for accurate cell division, with key roles played by kinetochore components CENP-E/kinesin-7 and Aurora B kinase. However, Aurora kinases both inhibit and promote congression, suggesting the presence of a larger signaling network. Our study demonstrates that centrosomes inhibit congression initiation when CENP-E is inactive by regulating the activity of kinetochore components. Depleting centrioles via Plk4 inhibition allows chromosomes near acentriolar poles to initiate congression independently of CENP-E. At centriolar poles high Aurora A enhances Aurora B activity, increasing phosphorylation of microtubule-binding proteins at kinetochores and preventing stable microtubule attachments in the absence of CENP-E. Conversely, inhibiting Aurora A or expressing a dephosphorylatable Hec1 mutant enables congression initiation without CENP-E. We propose a self-limiting feedback mechanism involving Aurora kinases and CENP-E that regulates the timing of chromosome movement by modulating kinetochore-microtubule attachments and fibrous corona expansion, with Aurora A activity gradient providing critical spatial cues for the network’s function.

## INTRODUCTION

Chromosome congression, the process by which chromosomes align at the spindle equator during early mitosis, is essential for accurate chromosome segregation and cell division^1,2^. The kinetochore motor protein CENP-E (Centromere-associated protein E) is recognized as a critical player in this process^2–6^. The prevailing view of CENP-E’s role during congression is that its plus-end directed motor activity moves chromosomes along microtubules to the spindle midplane, independently of their biorientation^7–10^. However, our recent observations challenge this understanding of CENP-E’s function in congression. We propose that chromosome congression is coupled to biorientation near the spindle poles through CENP-E, which stabilizes end-on attachments at polar kinetochores via its interaction with BubR1^11^. Thus, chromosome congression requires conversion from lateral to end-on attachments^12–14^. Notably, CENP-E is essential only for the congression of chromosomes near centrosomes^8,15^, and biorientation in unperturbed spindles occurs only outside the centrosome-proximal regions^16,17^. Yet, the role of centrosome signaling in both chromosome biorientation and congression is unclear.

Centrosomes are non-membranous organelles involved in various cellular processes, including cell division and ciliogenesis^18^. Majority of somatic cells in vertebrates contain two centrosomes, each centered around a pair of centrioles. These cylindrical structures, composed of microtubules arranged in a nine-fold symmetry, anchor and organize the pericentriolar material, which is crucial for microtubule nucleation and spindle formation^18^. Previous studies have demonstrated that centrioles are not essential for chromosome segregation in mitotic vertebrate cells^19,20^, and in oocytes which naturally lack centrioles^21^. Centrosome-associated kinases such as Aurora A are critical for centrosome maturation, which involves enlargement, increased microtubule nucleation capacity, and the establishment of spindle poles prior to mitosis^22^. During congression movement, Aurora kinases activate CENP-E through phosphorylation, thereby promoting chromosome congression in the vicinity of centrosomes^10,23^. However, in the accompanying manuscript we have shown that Aurora kinases also inhibit congression initiation when CENP-E is inactive of absent^11^. These seemingly paradoxical activities of Aurora kinases within the same process suggest the existence of signaling feedback among different components of the mitotic spindle. However, the nature of this feedback and its regulation remain unknown.

In this study, we investigated the role of centrosomes in regulating chromosome congression under varying levels of CENP-E activity by combining RNA interference (RNAi), expression of phospho-mutants, acute chemical inhibition, phospho-specific antibodies, and large-scale live-cell imaging approaches. Our findings reveal that centrioles, through the activity of Aurora A kinase, inhibit the congression of polar chromosomes when CENP-E is non-functional. By depleting centrioles through Polo-like kinase 4 (Plk4) inhibition, we show that acentriolar poles enable chromosomes to initiate congression independently of CENP-E, in contrast to centriolar poles. The inhibitory effect of centrioles on congression initiation is mediated by high Aurora A activity at centriolar poles. Close to centrosomes Aurora A overactivates Aurora B at kinetochores leading to increased phosphorylation of the KMN network (comprising the Knl1 complex, the Mis12 complex, and the Ndc80 complex), which in turn prevents stable microtubule attachments at kinetochores. Finally, expression of constitutively dephosphorylated Hec1 is sufficient to induce congression movement in the absence of CENP-E if fibrous corona disassembles. We thus propose a negative feedback loop in which Aurora B kinase, activated by Aurora A near centrosomes, destabilizes end-on attachments at the spindle poles and promotes fibrous corona expansion, while CENP-E, also activated by Aurora kinases, counteracts this destabilization to promote chromosome congression. These insights offer a new understanding of the complex molecular interplay between centrosomes and kinetochores during mitosis.

## RESULTS

### Centrioles delay the initiation of congression when CENP-E is inactive

Chromosome congression is notably delayed in cells lacking CENP-E, with polar chromosomes often lingering near the centrosomes before initiating movement toward spindle equator^8,11,24^. Based on this, we hypothesized that centrosomes might inhibit the initiation of congression of polar chromosomes when CENP-E is inactive or absent. To test this hypothesis, we imaged non-transformed human RPE-1 cells expressing CENP-A-GFP and centrin1-GFP^25^ by confocal microscopy after acute reactivation of CENP-E and after depletion of CENP-E by RNA interference (Extended Data Fig. 1a and b)^11^. We then tracked the movement of polar kinetochores over time (see Methods) and measured the duration of congression as a proxy of the likelihood of congression initiation for each pair of sister kinetochores with respect to their initial distance from the nearest centrosome. From this analysis, we found that the likelihood of congression initiation increases with the distance of the kinetochore pair from the centrosome, prominently in the absence of CENP-E and weakly after its acute reactivation (Extended Data Fig. 1c and d). This suggests that centrosomes play an inhibitory role in the initiation of congression in the absence of CENP-E.

To investigate the inhibitory role of centrosomes in chromosome congression in more detail, we employed a targeted imaging assay designed to systematically explore the interplay between key molecular players at kinetochores and centrosomes. Using the advanced lattice light-sheet microscopy approach described in the accompanying manuscript^11^, we captured high-resolution time-lapse images of live mitotic RPE-1 cells with labelled centromeres and centrosomes. We combined this imaging approach with CENP-E perturbation techniques that allowed us to vary CENP-E activity: CENP-E depletion, CENP-E inhibition, and control DMSO-treated cells^11^ (Fig. 1a). If centrosomes inhibit the congression of polar chromosomes in the absence of CENP-E, as we hypothesized, then depleting centrioles might allow these chromosomes to initiate congression independently of CENP-E. To deplete centrioles, we inhibited polo-like kinase 4 (Plk4) activity using centrinone, a small molecule inhibitor that blocks the centriole duplication cycle^19^. By varying the duration of treatment, we generated spindles with different numbers of centrioles at each pole—specifically 2:2 without treatment, 1:1 after one day, 1:0 after two days, and 0:0 after more than three days of treatment—and combined these with CENP-E perturbations (Fig. 1a and b).

**Fig. 1.**
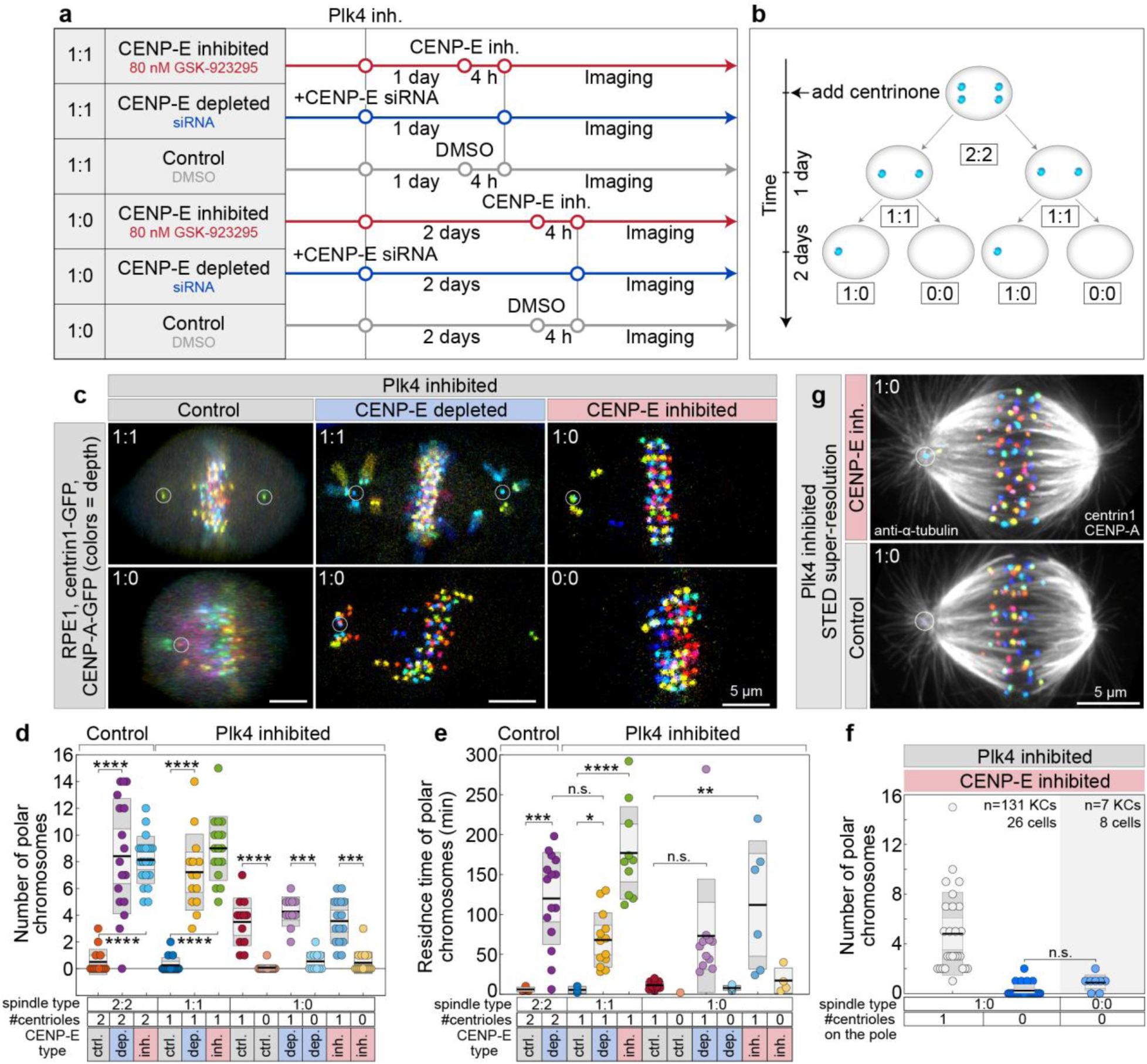
Centrioles inhibit congression initiation without CENP-E activity. **(a)** Protocol schematic to modulate CENP-E activity alongside variable centriole numbers per spindle pole obtained by continuous Plk4 inhibition. **(b)** Diagram showing progressive centriole depletion after continuous Plk4 inhibition by 300 nM centrinone. **(c)** Representative live images of RPE-1 cells expressing CENP-A-GFP and centrin1-GFP (color-coded by depth) with varying centriole counts (white circles), 5 minutes after end of prometaphase spindle elongation, under indicated treatments. **(d, e)** Quantification of polar chromosome number (d) and residence time at spindle poles (e) after prometaphase spindle elongation, comparing cells with different centriole numbers. Category “1” includes centriolar poles from both 1:1 and 1:0 spindles. Black lines indicate means; gray areas show 95% confidence intervals and standard deviations. Number of cells per condition: 14, 15, 22, 16, 14, 18, 16, 16, 12, 12, 16, 16 (d, e), each from ≥3 replicates. **(f)** Number of polar chromosomes after treatments in cells with varying centriole numbers, fixed prior to measurement. Pooled data from more than ≥3 replicates. **(g)** STED microscopy images of RPE-1 cells immunostained for α-tubulin (grey), expressing CENP-A-GFP and centrin1-GFP (color-coded by depth). All images are maximum projections. Statistics: ANOVA with Tukey post-hoc test. Symbols indicate: n.s., P > 0.05; *, P ≤ 0.05; **, P ≤ 0.01; ***, P ≤ 0.001; ****, P ≤ 0.0001; inh., inhibited; depl., depleted; siRNA, small interfering RNA.

As a proxy for congression efficiency, we quantified the number of polar chromosomes 5 minutes after spindle elongation, the residence time of polar chromosomes at the spindle poles, and the total duration of mitosis (Methods). Following CENP-E perturbations, spindles with a 1:1 or 2:2 centriole configuration had an average of about eight polar chromosomes per spindle, compared to fewer than one chromosome under control conditions in both spindle types (Fig. 1c and d; and Video 1). In 1:0 spindles following CENP-E perturbations, the centriolar poles (labeled “1”) contained approximately four polar chromosomes, similar to their counterparts in 1:1 spindles (Fig. 1c, d; and Video 1). The occurrence of polar chromosomes at centriolar poles in 1:0 spindles was independent of CENP-E activity (Fig. 1d). The acentriolar poles (labeled “0”) in 1:0 spindles consistently had an average of around one polar chromosome, regardless of CENP-E activity (Fig. 1c and d; and Video 1). Importantly, polar chromosomes persisted significantly longer at centriolar poles than at acentriolar poles across all spindle types when CENP-E was perturbed (Fig. 1e), resulting in a marked prolongation of mitosis (Extended Data Fig. 1e). Furthermore, completely acentriolar 0:0 spindles, which were rarely observed due to p53-dependent arrest in RPE-1 cells^19^, exhibited a low number of polar chromosomes under CENP-E inhibition, similar to the acentriolar poles in 1:0 spindles (Fig. 1c and f; and Video 1). These findings suggest that centrioles specifically limit the initiation of polar chromosome congression when CENP-E activity is compromised.

The low number of polar chromosomes observed at acentriolar poles may be explained by the transient monopolarization of 1:0 spindles^26^. However, we observed that forced transient monopolarization through Eg5 inhibitor washout, followed by CENP-E inhibition, did not increase the number of polar chromosomes compared to untreated 2:2 spindles (Extended Data Fig. 2a)^11^. This suggests that the inhibitory effect of centrioles on polar chromosome congression in 1:0 spindles without CENP-E is independent of passage through the transient monopolar state. To observe differences in microtubule organization between centriolar and acentriolar poles we imaged microtubules using STED microscopy. However, we observed no significant differences in microtubule organization between centriolar and acentriolar poles, aside from the absence of astral microtubules at the acentriolar pole (Fig. 1g). These findings suggest that in non-transformed cells, the ability to initiate chromosome congression when CENP-E is inactive depends on the distance of the chromosome to the centrosome.

### Centrioles regulate congression initiation without affecting congression dynamics

To determine whether the inhibitory effect of centrioles on congression is dependent on chronic Plk4 inhibition, we investigated whether centriole dislocation from spindle pole during prolonged mitosis could induce the congression of polar chromosomes independently of the long-term effects of Plk4 inhibition. We focused on rare instances (<10%, n=5 out of 55 cells) where spontaneous acute dislocation of a centriole from the spindle pole occurred in 1:1 spindles under CENP-E depletion in RPE-1 cells (Extended Data Fig. 2b), a phenomena observed previously in cells undergoing prolonged mitosis^27,28^. Fascinatingly, in all observed cases, polar chromosomes initiated congression almost immediately following the displacement of a centriole from the centrosome in CENP-E inhibited cells (Extended Data Fig. 2b). This finding suggests that the mere presence of centrioles limits the congression of polar chromosomes in the absence of CENP-E.

Having found that centrioles inhibit congression initiation in the absence of CENP-E, we next asked whether centriole number influences the dynamics of congression movement. Congression velocity and interkinetochore distance were indistinguishable among 1:1, 1:0, and 2:2 spindles following CENP-E inhibition in RPE-1 cells (Fig. 2a; Extended Data Fig. 2c–g)^11^. These results suggest that centrioles primarily inhibit the initiation of congression, but not the subsequent movement of polar chromosomes toward the metaphase plate.

**Fig. 2.**
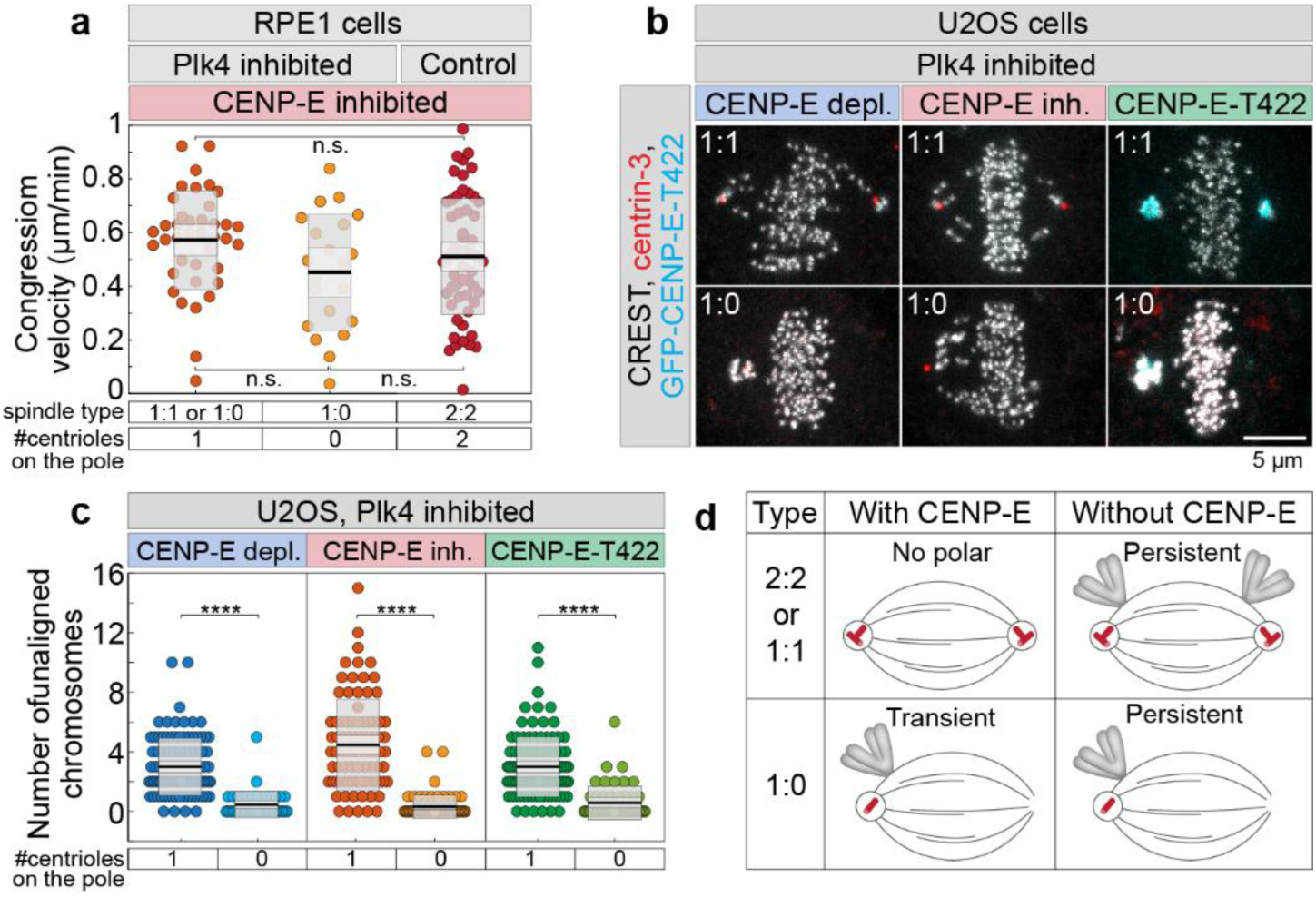
Centrioles inhibit congression initiation upon phospho-null CENP-E expression without affecting congression velocity. (**a**) Velocity of chromosome congression movement in RPE1 cells during the 6-minute period that precedes complete alignment for each initially polar kinetochore pair in different treatments and number of centrioles. (**b**) Representative examples of fixed U2OS cells expressing inducible GFP-CENP-E-T422 (cyan) and immunostained for human centromere protein (CREST, grey) and centrin-3 (red), with different numbers of centrioles on each pole (rows) and for different treatments (columns) as indicated on top. (**c**) Number of polar chromosomes after different treatments of U2OS cells, as indicated on top, and with different numbers of centrioles, as indicated on bottom. (**d**) Schemes depicting the summary of experimental results for the number and residence time of polar chromosomes on spindle poles for cells characterized by different numbers of centrioles with or without CENP-E activity. All images are maximum projections. Numbers: 125 kinetochores from 36 cells (a), and 225 cells (c), each from ≥2 replicates. Statistics: ANOVA with post-hoc Tukey test. Symbols indicate: ****, P ≤ 0.0001; inh., inhibited; depl., depleted; siRNA, small interfering RNA.

To test whether selective disruption of CENP-E activity, without complete depletion or inducing rigor microtubule binding of the motor, would also bias accumulation of polar chromosomes to the centriolar pole, we used osteosarcoma U2OS cells engineered for doxycycline-inducible expression of phospho-null Threonine 422 (T422A) mutant lacking the Aurora A/B-specific phosphorylation site^23^. To compare different modes of CENP-E perturbation in transformed cells, we analyzed three conditions: 1) endogenous CENP-E depletion, 2) pharmacological inhibition of CENP-E, 3) and expression of T422A CENP-E following depletion of endogenous CENP-E. Immunofluorescence microscopy was performed under continuous centrinone treatment for 2 days to generate mixed populations of cells with either centriolar (1:1) or mixed centriolar and acentriolar (1:0) spindle poles (Fig. 2b). We quantified the number of polar chromosomes relative to the number of centrioles on the spindle pole. Consistent with our findings in RPE-1 cells (Fig. 1e, f), polar chromosomes predominantly accumulated at centriolar poles under all conditions that impaired CENP-E activity, including expression of the T422A mutant (Fig. 2c). These results support a model in which CENP-E-dependent congression of polar chromosomes is spatially biased toward centriolar spindle poles, both in non-transformed and transformed cells and across distinct modes of CENP-E perturbation (Fig. 2d).

### Hyperactivated Aurora B near centrosomes enhances KMN phosphorylation and delays congression without CENP-E

What molecular factors delay congression initiation near centrioles in the absence of CENP-E? Since we have shown that inhibition of Aurora A kinase induces the congression of polar chromosomes in the absence of CENP-E or its activity^11^, we hypothesized that centrosomal Aurora A activity inhibits chromosome congression at centriolar poles when CENP-E is absent. To test this hypothesis, we quantified the levels of active Aurora A (pAurA, pT288-Aurora A) on spindle poles with varying numbers of centrioles after continuous Plk4 inhibition in RPE-1 cells, under both control and CENP-E inhibited conditions. Intriguingly, we found that acentriolar poles exhibited a 4-fold lower level of pAurA compared to centriolar poles, whereas the maximum difference between the two poles in 1:1 spindles was on average 10% (Fig. 3a and b). The observed differences in pAurA levels between poles were not dependent on CENP-E (Fig. 3a and b), suggesting that CENP-E does not regulate the activity of Aurora A. These findings, combined with our observation that Aurora A inhibition promotes congression in the absence of CENP-E^11^, indicate that polar chromosomes rely on CENP-E activity for congression due to the high activity of Aurora A near centrosomes.

**Fig. 3.**
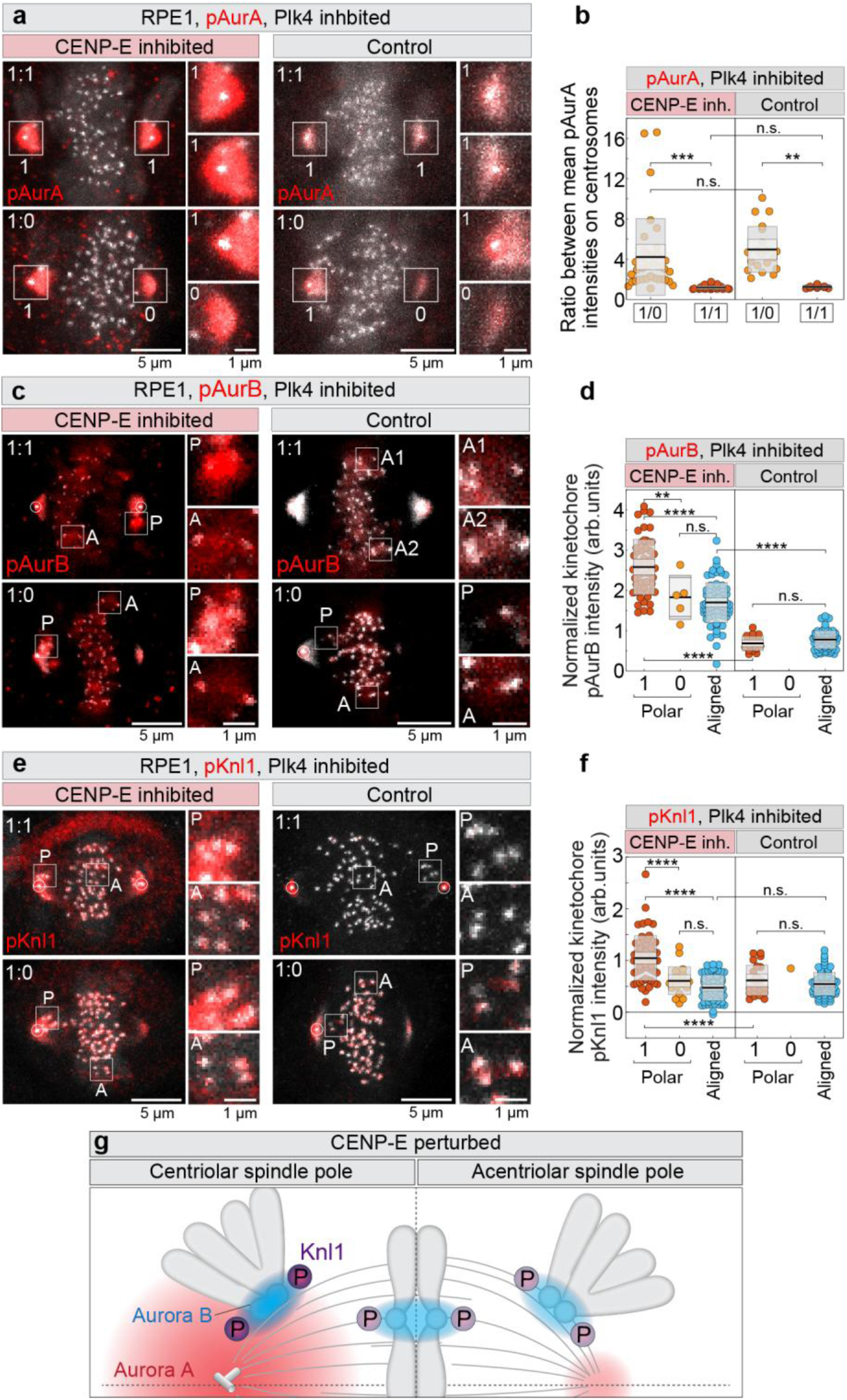
Aurora B activity on kinetochores near centrioles is enhanced causing hyperphosphorylation of the KMN network when CENP-E is inactive. (**a**, **c**, **e**) Representative RPE-1 cells expressing CENP-A-GFP and centrin1-GFP (grey) with centrioles marked by white circles, immunostained for phosphorylated residues of Aurora A (pT288, a), Aurora B (pT232, c), and Knl1 (pS24, pS60, e) (red) after indicated treatments and centriole numbers. Images are maximum projections with enlarged insets of polar (P) and aligned (A) kinetochores on the right. (**b**, **d**, **f**) Quantification of the ratio of phosphorylated Aurora A intensity at centriolar versus acentriolar spindle poles (b), and normalized phosphorylation levels of Aurora B (d) and Knl1 (f) under different treatments and centriole counts. (**g**) Schematic representation of kinetochore phosphorylation levels in the absence of CENP-E activity. Aurora B activity is depicted as a blue gradient centered on the centromere, with intensity scaling according to phosphorylation level. Knl1 phosphorylation (P) is represented by the intensity of purple circles. The spindle features one centriolar pole (left; centrioles shown schematically) and one acentriolar pole (right). The Aurora A activity gradient (red) is stronger and extends further from the centriolar pole. Phosphorylation patterns are illustrated for both polar and aligned chromosomes. Numbers: 88 cells (b), 179 kinetochores from 42 cells (d), and 227 kinetochores from 63 cells (f), each from ≥2 replicates. Statistics: ANOVA with post-hoc Tukey’s HSD test. Symbols indicate: n.s., P > 0.05; **, P ≤ 0.01; ***, P ≤ 0.001; ****, P ≤ 0.0001; inh., inhibited; depl., depleted; KC, kinetochore; NT, non-targeting; arb., arbitrary.

We hypothesized that the high Aurora A activity at centriolar spindle poles might lead to increased activation of kinetochore-localized Aurora B kinase on polar chromosomes. High Aurora B activity would affect the phosphorylation of its downstream targets, such as Knl1, which is the component of the KMN network^29^, thus blocking congression. Indeed, we have observed that the mean signals of pAurB (pT232-Aurora B) and pKnl1 (pS24, pS60-Knl1)^30^ on polar kinetochores near centriolar poles were significantly higher compared to aligned kinetochores and the rare polar kinetochores at acentriolar poles when CENP-E was inhibited (Fig. 3c-f; and Extended Data Fig. 3a and b).

To confirm the specificity of phospho-antibodies as indicators of Aurora B activity, we performed acute Aurora B inhibition by 3 μM ZM-4473 and assessed the localization and signal intensity of pAurB and pKnl1. Both signals were almost completely lost from kinetochores following inhibition, except for a small region near the centrioles, confirming their specificity as markers of Aurora B activity (Extended Data Fig. 3c). In contrast, the intensity and localization of pHec1 (pS55-Hec1)^31,32^ remained unchanged following acute Aurora B inhibition (Extended Data Fig. 3c), suggesting that S55 phosphorylation does not specifically reflect Aurora B activity, consistent with a recent report^33^. This likely explains the uniform pHec1 levels at polar kinetochores regardless of the presence of centrioles on spindle poles and CENP-E activity (Extended Data Fig. 3d, e), which contrasts with the spatial variation and CENP-E dependence observed for pKnl1 and pAurB (Fig. 3c–f). In DMSO-treated control cells, pAurB and pKnl1 levels were comparable between aligned and rare polar kinetochores and were significantly lower than those observed in the same kinetochore groups under CENP-E inhibition (Fig. 3c–f).

These results suggest that in the absence of CENP-E activity, overactivated Aurora A at centriolar poles enhances the activity of kinetochore-localized Aurora B. Overactivated Aurora B increases the phosphorylation of downstream outer kinetochore proteins, such as Knl1, and probably KMN network in general (Fig. 3g). Hyperphosphorylated KMN inhibits the stabilization of end-on microtubule attachment to kinetochores, thereby preventing chromosome congression.

### Aurora A overactivates Aurora B on kinetochores near centrosomes

Given that polar kinetochores near centrosomes in the absence of CENP-E show elevated Aurora B phosphorylation, we hypothesized that Aurora A at spindle poles directly promotes Aurora B activity at kinetochores, consistent with their shared substrate recognition motifs^34–36^. To test this, we first inhibited CENP-E and then acutely inhibited Aurora A by the highly specific inhibitor MLN8054^37^, using a range of concentrations and treatment durations. We then compared pAurB levels at kinetochores in CENP-E-inhibited cells following acute treatment with Aurora A inhibitor or DMSO (Fig. 4a). Acute inhibition of Aurora A significantly reduced Aurora B activity on polar kinetochores, bringing it to levels observed on aligned kinetochores in CENP-E–inhibited cells (Fig. 4a–c). To validate this finding, we tested two additional Aurora A–specific inhibitors: TCS7010^38^, which also reduced pAurB asymmetry on polar versus aligned kinetochores similar to MLN8054, and Alisertib^39^, which had no significant impact under the tested conditions (Fig. 4d, e). Consistent with these results, only MLN8054 and TCS7010 caused a ∼2-fold decrease in the number of polar chromosomes and a significant reduction in spindle length^40^ compared to CENP-E inhibition alone, without altering pAurB levels at aligned kinetochores (Fig. 4f, g; Extended Data Fig. 3f). These findings support a model in which Aurora A directly enhances Aurora B activity at polar kinetochores when CENP-E is inactive.

**Fig. 4.**
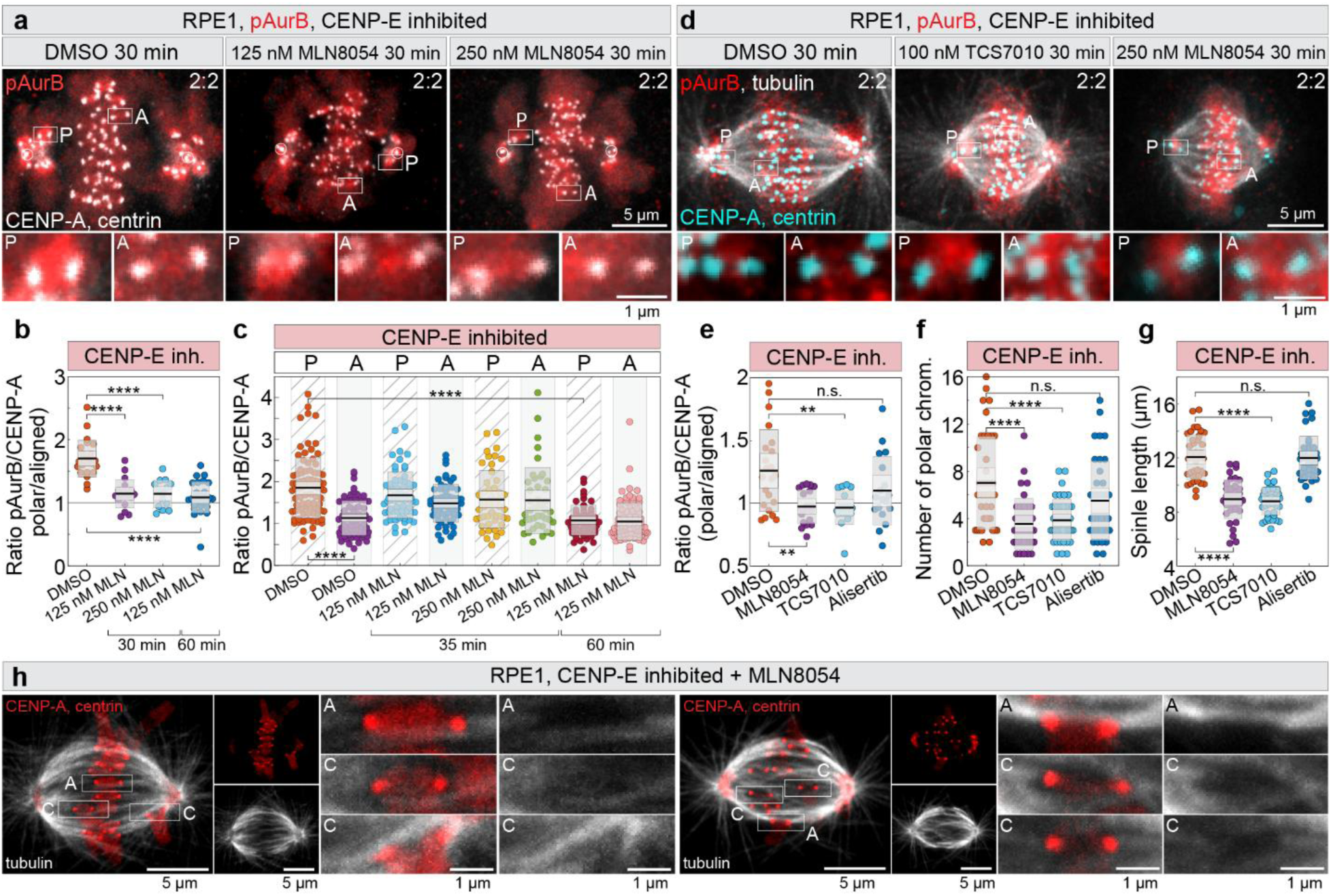
Aurora A on centrosomes upregulates Aurora B activity on kinetochores. (**a**) Representative RPE-1 cells expressing CENP-A-GFP and centrin1-GFP (grey) with centrioles marked by white circles, after indicated treatments, showing polar (P) and aligned (A) kinetochores immunostained for pT232-Aurora B (red). (**b**) Ratio of pT232-Aurora B intensity normalized to CENP-A on polar vs. aligned kinetochores per cell under indicated treatments. MLN, MLN8237. (**c**) Average pT232-Aurora B levels normalized to CENP-A on all kinetochore groups after indicated treatments. (**d**) Representative cells expressing CENP-A-GFP and centrin1-GFP (cyan) immunostained for pT232-Aurora B (red) and α-tubulin (grey). (**e**–**g**) Quantification of pT232-Aurora B intensity ratio on polar vs. aligned kinetochores (e), number of polar chromosomes (f), and spindle length (g) after treatments. (**h**) Two RPE-1 cells imaged by super-resolution Airyscan microscopy showing congressing (C) and aligned (A) kinetochores with α-tubulin (grey) and CENP-A-GFP and centrin-GFP (red). Images are deconvolved projections of 2–5 z-planes. Numbers: 571 kinetochores in 91 cells (b, c), 436 kinetochores from 76 cells (e-g), all from ≥2 replicates. Statistics: ANOVA with post-hoc Tukey’s HSD test. Symbols indicate: n.s., P > 0.05; **, P ≤ 0.01; ***, P ≤ 0.001; ****, P ≤ 0.0001; KC, kinetochore.

Our model suggests that downregulation of Aurora kinases promotes chromosome congression by stabilizing end-on attachments at polar kinetochores. To test this, we stained tubulin in cells pre-treated with a CENP-E inhibitor, followed by acute treatment with the Aurora A inhibitor MLN8054 (Fig. 4h). Super-resolution Airyscan imaging^41^ revealed that most kinetochores located between the centrosome and the metaphase plate had end-on microtubule attachments after acute MLN8054 treatment in CENP-E–inhibited cells (n=15 cells) (Fig. 4h, Extended Data Fig. 3g). These results are consistent with our recent findings showing that acute CENP-E reactivation leads to stabilized end-on attachments on congressing chromosomes^11^. Together, these findings suggest that Aurora A modulates Aurora B activity near spindle poles, reinforcing our model of localized regulation of chromosome congression.

If Aurora A at the centrosomes enhances Aurora B activity in the absence of CENP-E, we would expect a decrease in Aurora B-mediated phosphorylation of outer kinetochore targets as chromosomes move away from the centrosome during congression. Indeed, we observed that the level of pKnl1 was inversely correlated with the distance of the kinetochore pair from the centrosome in the absence of CENP-E activity (Extended Data Fig. 3h). In conclusion, chromosome congression is accompanied by a reduction in Aurora B activity on kinetochores, leading to decreased phosphorylation of KMN network components. The reduction of Aurora B activity on congressing chromosomes is crucial for the stable attachment of microtubules to kinetochores^42^ and is significantly accelerated by the motor activity of CENP-E.

### Constitutive Hec1 dephosphorylation is sufficient to induce congression without CENP-E

To test if downregulation of Aurora kinase-mediated Hec1 phosphorylation at polar kinetochores is sufficient to initiate chromosome congression, we used HeLa cells engineered to express an inducible, siRNA-resistant GFP-Hec1-9A phopho-mutant^43^ (Fig. 5a, top, b). This mutant mimics the constitutive dephosphorylation of Hec1 at nine phosphorylation sites, which are phosphorylated by Aurora B kinase^44^. Expression of Hec1-9A in the presence of endogenous Hec1 mildly affected chromosome congression but caused an increase in interkinetochore distance of aligned kinetochore pairs (Fig. 5b, c; Extended Data Fig. 3i), consistent with previous reports^43,44^. Chromosome congression was completely disrupted by depletion or inhibition of CENP-E, as well as Hec1 depletion, leading to an accumulation of polar chromosomes, consistent with previous studies (Fig. 5b-d)^8,45^. Interestingly, both depletion and inhibition of CENP-E caused severe congression defects in cells expressing endogenous Hec1, even upon Hec1-9A overexpression (Fig. 5c). This suggests that Hec1-9A overexpression cannot compensate for CENP-E disruption in the presence of endogenous Hec1, despite its dominant-negative effect on interkinetochore tension and SAC satisfaction^43,46^.

**Fig. 5.**
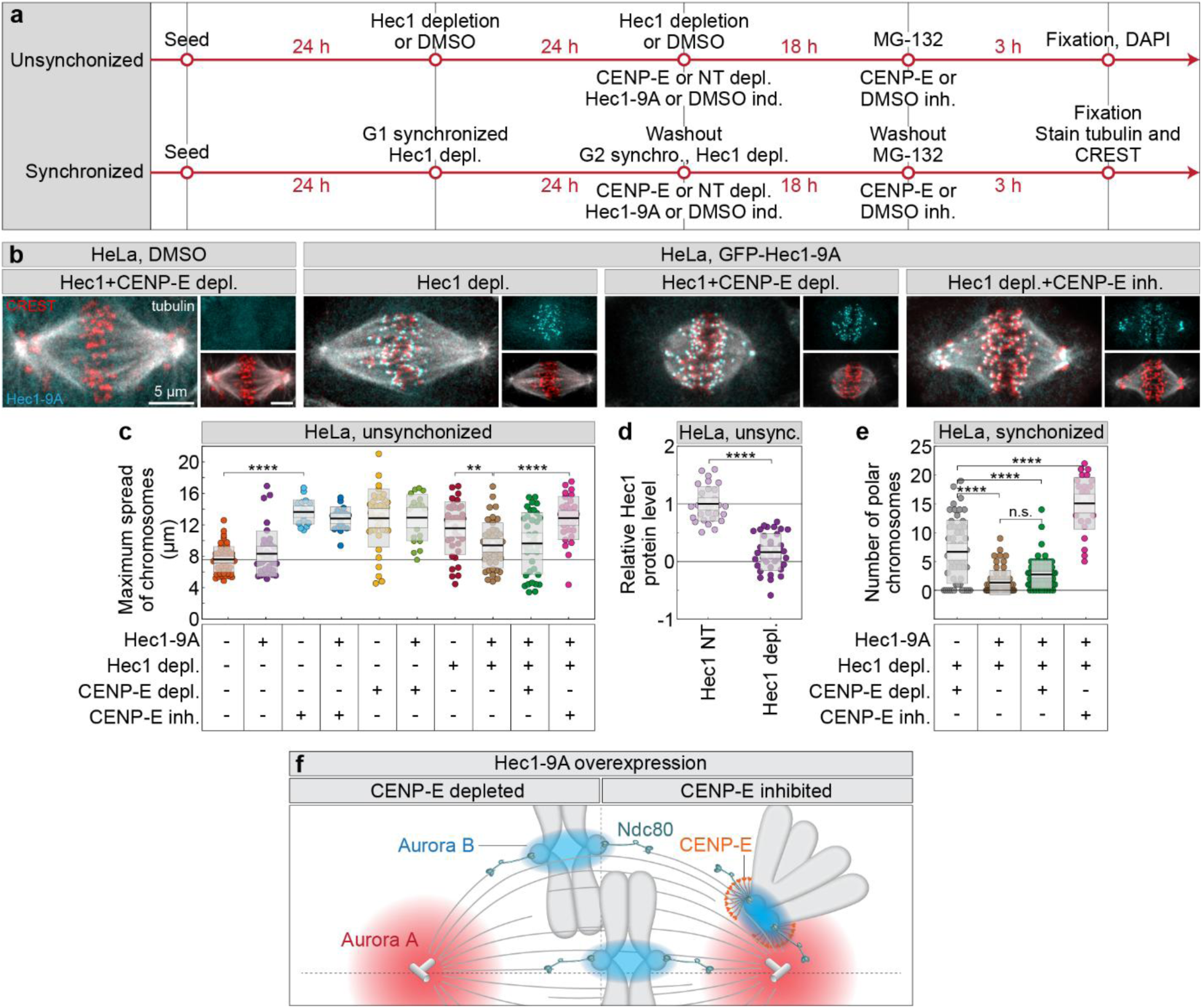
Aurora kinase–mediated Hec1 phosphorylation impairs chromosome congression in the absence of CENP-E. (**a**) Schematics of two experimental workflows using either unsynchronized HeLa cells (top) or cells synchronized in G1 and G2 during protein depletion (bottom), with treatments above the line common to all conditions and those below subgroup-specific. (**b**) Representative images of HeLa cells synchronized in mitosis with MG-132, without (left) or with (all others) GFP-Hec1-9A expression (cyan), immunostained for α-tubulin (gray) and centromeres (CREST, red) in treatments indicated on top. (**c**) Maximum chromosome spread in unsynchronized cells after indicated treatments. (**d**) Hec1 levels measured within the DAPI-stained DNA region after Hec1 3’UTR depletion, normalized to non-targeting controls. (**e**) Number of polar chromosomes in synchronized HeLa cells after indicated treatments. (**f**) Schematic representation of chromosome congression capacity in cells overexpressing Hec1-9A with endogenous Hec1 removed. In CENP-E–depleted cells (left pair), chromosome congression is efficient, and the Ndc80 complex forms stable end-on attachments with microtubules—similar to aligned kinetochores (bottom pair). In contrast, under CENP-E inhibition (right pair), the fibrous corona (simplified here as CENP-E) prevents stabilization of end-on attachments, blocking Hec1– microtubule interaction and impairing congression at polar kinetochores. Aurora B activity is shown as a blue gradient centered on the centromere, while Aurora A is represented as a red gradient centered on a pair of schematically depicted centrioles. Numbers: 571 kinetochore pairs from 91 cells (c), 327 cells (d) 203 cells (e), all from ≥3 replicates. Statistics: ANOVA with post-hoc Tukey’s HSD test. Symbols indicate: n.s., P > 0.05; **, P ≤ 0.01; ***, P ≤ 0.001; ****, P ≤ 0.0001; inh., inhibited; depl., depleted; NT, non-targeting; KC, kinetochore.

Based on our finding that acute inhibition of Aurora kinases induces chromosome congression independently of CENP-E^11^, we hypothesized that constitutive dephosphorylation of Hec1 might compensate for disruptions of CENP-E function after depletion of endogenous Hec1. To control for potential mitotic prolongation effects following combined Hec1 and CENP-E perturbations, we synchronized cells in G1 before depleting endogenous Hec1, CENP-E, or both, thereby ensuring depletion prior to mitotic entry. After release into G2 and mitosis, cells were arrested in metaphase using a proteasome inhibitor for a maximum of 2 hours (Fig. 5a, bottom).

We assessed chromosome congression efficiency by immunostaining cells for tubulin and centromeres under four conditions, all in the context of endogenous Hec1 depletion: and (1) CENP-E depletion, (2) Hec1-9A expression, (3) Hec1-9A expression with CENP-E depletion, (4) Hec1-9A expression with CENP-E inhibition. Expression of Hec1-9A in cells lacking endogenous Hec1 rescued major chromosome congression defects in both synchronized and unsynchronized cells (Fig. 5b, c, e), as previously reported^43,44^. Remarkably, expression of Hec1-9A also significantly improved chromosome congression in cells depleted of both endogenous Hec1 and CENP-E (Fig. 5b). The improvement was reflected in both the maximum chromosome spread and the average number of polar chromosomes, which closely matched those observed in cells with intact CENP-E, in synchronized (Fig. 5b, e, Extended data Fig. 3j) as well as unsynchronized cells (Fig. 5c, Extended data Fig. 3i). In all conditions where Hec1-9A was overexpressed, aligned chromosomes had a large interkinetochore distance and showed end-on microtubule attachments (Fig. 5b). Surprisingly, Hec1-9A overexpression did not rescue the congression defects caused by CENP-E inhibition in either synchronized or unsynchronized cells (Fig. 5b, c and e). This is likely due to hyperexpansion of the fibrous corona on polar kinetochores following CENP-E inhibition, which interferes with the stabilization of microtubule attachments^11^. Consistent with this, the location and interkinetochore distance of polar kinetochores in cells overexpressing Hec1-9A under CENP-E inhibition (Fig. 5b) were similar to those under CENP-E inhibition alone (Fig. 4d). These findings suggest that constitutive Hec1 dephosphorylation is sufficient to drive chromosome congression in the absence of CENP-E and endogenous Hec1, but only when excessive fibrous corona expansion is avoided.

## DISCUSSION

In this study, we propose that the timing of chromosome movement is regulated by the interplay between activity of the CENP-E molecular motor at kinetochores and the chromosomes’ proximity to centrosomes, which determines their exposure to the Aurora A gradient (Fig. 6). Previous research has shown that Aurora kinases phosphorylate and activate CENP-E near centrosomes^10,23^. The findings presented in the accompanying manuscript^11^, together with the results shown here, demonstrate that Aurora kinases inhibit the initiation of chromosome congression in the absence of CENP-E. To reconcile these seemingly conflicting roles of Aurora kinases within the same process, we propose a feedback loop where kinetochore activity is influenced by the proximity to centrosomes, where the Aurora A gradient is concentrated in somatic human cells (Fig. 6)^47^. Without CENP-E, Aurora A overactivates Aurora B and its targets, such as KMN complex^13,30,42,44^ and the fibrous corona components^48,49^, inhibiting end-on attachment stabilization and congression^8,11,50^ (Fig. 6). However, direct biochemical evidence for an Aurora A–Aurora B interaction has yet to be established. Aurora A also directly phosphorylates KMN components, including Hec1, near centrioles^32,50^. The specific contributions of Aurora kinases to Hec1 phosphorylation sites (S44, S55, S69 and others) and role of distinct phosphorylation sites in congression inhibition remain unclear^33,51^. Future studies will hopefully clarify this.

**Fig. 6.**
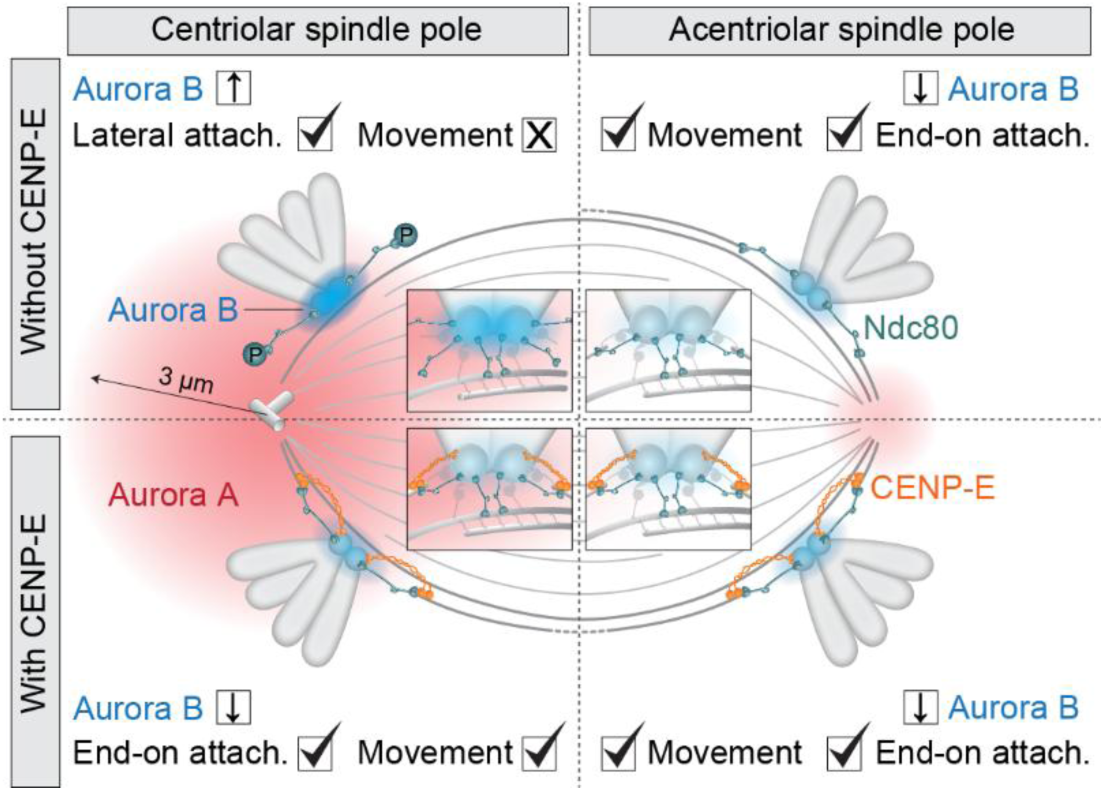
Spatial regulation of chromosome congression via an Aurora B–CENP-E feedback loop controlled by proximity to Aurora. **A.** The initiation of end-on kinetochore-microtubule attachments and chromosome congression depends on the distance between chromosomes and the centrosome. In the absence of CENP-E (represented as orange motor), polar chromosomes near centriolar centrosomes (top left) remain laterally attached to microtubules due to high Aurora A (AurA, represented as red gradient with a radius of ∼3 µm) activity, which enhances Aurora B (AurB, represented as blue gradient on centromeres and kinetochores) activation leading to hyperphosphorylation of outer kinetochore proteins (represented by Ndc80 complex, blue) and expansion of fibrous corona (not shown), preventing stable end-on attachments. When CENP-E is present (bottom left), Aurora B activity at the kinetochore is reduced, allowing end-on conversion, biorientation, and congression, aided by BuBR1 (not shown). Without centrosomes and CENP-E (top right), the absence of the AurA gradient lowers Aurora B activity, permitting end-on conversion and congression independently of CENP-E. In the presence of CENP-E near acentriolar poles (bottom right), chromosome congression proceeds similarly to centriolar poles with CENP-E. Insets show kinetochore-level detail per condition. Symbols indicate: X = absence or low activity; ↓ = downregulation; ✓ = presence; attach. = attachment.

Our findings support a model in which kinetochore-centrosome feedback is not solely driven by the attachment status of microtubules to kinetochores but involves a bidirectional regulatory loop between Aurora kinases and CENP-E. While CENP-E promotes stable end-on attachments that suppress Aurora B activity, Aurora A and B themselves increase CENP-E activity and localization^10,23^. This interdependence suggests that changes in kinetochore signaling reflect not just passive responses to the attachment state but active feedback between motor activity and kinase signaling near the spindle poles. When CENP-E is present on kinetochore, it reduces Aurora B activity, promoting end-on attachments, fibrous corona removal and effective congression initiation (Fig. 6)^11^. If the Aurora A gradient is weakened, as in acentriolar spindle poles or after acute Aurora A inhibition, Aurora B activation decreases, allowing end-on attachment and CENP-E-independent congression of polar chromosomes (Fig. 6). However, the weak activity of both Aurora kinases is associated with more frequent formation of aberrant attachments of kinetochores to microtubules close to centrosomes, increasing chromosome mis-segregation rates during anaphase^19,50,52^.

During unperturbed mitosis in healthy human cells, chromosomes typically remain at least 3 μm away from the centrosome and achieve rapid biorientation outside this region^16,25,53^. We propose that in cells with active CENP-E most chromosomes rapidly establish end-on attachments and become bioriented close to the spindle surface by capturing microtubules extending from the opposite spindle half^16,54^. Thus, while CENP-E supports microtubule stabilization at all kinetochores, as indicated by reduced microtubule density in kinetochore fibers upon its perturbation^28^, its loss predominantly affects polar chromosomes without causing widespread detachment of those positioned farther from centrosomes.

The model we present clarifies the signaling dynamics between centrosomes and kinetochores during the initiation of chromosome congression. It explains why pole-proximal kinetochores located within the Aurora A gradient rely on CENP-E for efficient congression during early mitosis^8^. This reliance aligns with findings that congression is not perturbed by constitutive Hec1 dephosphorylation or acute Aurora kinase inhibition^11^, as the KMN network retains maximum microtubule affinity under this conditions^43,44^. Recent findings also show that kinetochores can modulate Aurora A activity via Mps1^55^, underscoring the feedback between centrosomes and kinetochores.

While our study highlights the central role of CENP-E and Aurora kinases in chromosome congression, other spindle-associated proteins are likely involved. HURP and CLASP proteins, for instance, stabilize and regulate kinetochore microtubules near chromosomes^56,57^, and the Ska complex enhances kinetochore–microtubule coupling under tension^58^. At the spindle poles and in their vicinity, factors such as TPX2, pericentriolar proteins, kinesin-13 depolymerases, the crosslinker NuMA, and the Augmin complex are all intricately linked to Aurora A signaling^36^. These pathways may act downstream of CENP-E and Aurora kinases to influence both congression efficiency and spatial bias, meriting further investigation.

Our findings suggest that it is not the presence of centrioles per se, but rather elevated Aurora A kinase activity that governs the feedback dynamics between spindle poles and kinetochores. In somatic cells, increased Aurora A activity is associated with canonical centrosomes, whereas oocytes maintain high Aurora A levels and form functional spindles without canonical centrosomes^59^. This may explain why oocytes, but not acentriolar poles in somatic cells, require CENP-E for chromosome congression despite the absence of centrioles^60,61^. We propose that spatial feedback mechanisms involving Aurora A may operate independently of centrioles, highlighting the need for further study of chromosome congression regulation in acentriolar contexts.

Regarding mitosis in diseased conditions, overexpression of Aurora A kinase and loss of the CHK2-BRCA1 tumor suppressor pathway, both of which lead to elevated Aurora A activity at centrosomes, are frequently observed in human cancers^62,63^. Increased Aurora A activity enhances microtubule assembly rates at kinetochores through mechanisms that are not yet fully understood, contributing to chromosomal instability in colorectal cancer cells^64^. Our findings suggest that elevated Aurora A activity could impede chromosome congression if not balanced by changes in other key components of the centrosome-kinetochore feedback loop, such as CENP-E/BubR1 and Aurora B kinase. Disruptions in this balance may explain the aberrant congression observed in colorectal cancer cells with overactive Aurora A kinase, which correlates with increased mitotic timing, and chromosome mis-segregation rates in those cells^64^.

Aurora A’s role as a major inhibitor of chromosome congression might also explain why chromosomes near the poles experience delayed congression, not only in the absence of CENP-E but also in other contexts where CENP-E is active. For example, in multipolar spindles with supernumerary centrosomes, chromosomes near coalesced spindle poles are often delayed in congression^65^, potentially due to their proximity to two centrosomes and their Aurora A gradients. Likewise, cancer cells from various origins frequently struggle with congressing pole-proximal chromosomes, leading to prolonged mitosis^66–68^. Although the cause of congression defects in cancer cells remains unclear, pole-proximal chromosomes are often mis-segregated into micronuclei, resulting in aneuploidy^66^. We propose that the chromosome congression defects observed across human tumors may result from imbalances in the signaling feedback that regulates chromosome congression, leading to disrupted stability of end-on attachments^33,69^.

### Limitations of study

It remains unclear why CENP-E inhibition was not effectively rescued by overexpressing Hec1-9A in the context of Hec1 depletion, unlike with CENP-E depletion. One possibility is that the Hec1-S69 phosphorylation site, absent in the Hec1-9A mutant^44^, is essential for bypassing the need for CENP-E under these conditions. Additionally, the extensive fibrous corona expansion might limit Hec1’s engagement with microtubules in the cells with inhibited CENP-E^49^. Further research is needed to clarify this. Moreover, Aurora A might have a more direct impact on KMN network components, potentially through specific phosphorylation sites on Hec1, such as S69, which were not examined in this study^32,50^.

In spindles where centrioles were removed by using the Plk4 inhibitor, potential indirect effects on chromosome congression cannot be fully excluded. For example, the asymmetric distribution of polar chromosomes may reflect the difference between the centriolar and acentriolar pole in their position and movement in addition to the difference in biorientation capacity. However, a similar trend was observed, albeit in a smaller number of cells, following spontaneous displacement of centrioles from spindle poles. Moreover, live imaging showed that chromosomes initially moved towards both poles, but they remained for a longer time at the centriolar pole (Video 1), supporting the robustness of our conclusions.

## ACKNOWLEDGEMENTS

We thank Alexey Khodjakov, Geert Kops and Marin Barišić for cell lines; Marin Barišić, Carlos Conde, Helder Maiato, and Andrea Musacchio for for reviewing and discussing the results; Julie Welburn for discussions and antibodies; Ivana Šarić for the drawings; and members of the Tolić group and Nenad Pavin group for constructive comments on the manuscript. This work was funded by the European Research Council (ERC Synergy Grant, GA Number 855158), the Croatian Science Foundation (HRZZ) through Swiss-Croatian Bilateral Projects (project IPCH-2022-10-9344), and projects co-financed by the Croatian Government and the European Union through the European Regional Development Fund—the Competitiveness and Cohesion Operational Programme: IPSted (Grant KK.01.1.1.04.0057) and QuantiXLie Center of Excellence (Grant KK.01.1.1.01.0004).

## AUTHOR CONTRIBUTIONS

K.V. and I.M.T conceived the project, K.V. performed all experiments, quantified, analyzed and presented the data, K.V. conceptualized and prepared original draft, K.V. and I.M.T. reviewed, edited and discussed the manuscript.

## MATERIALS & METHODS

### Reagents and Tools table

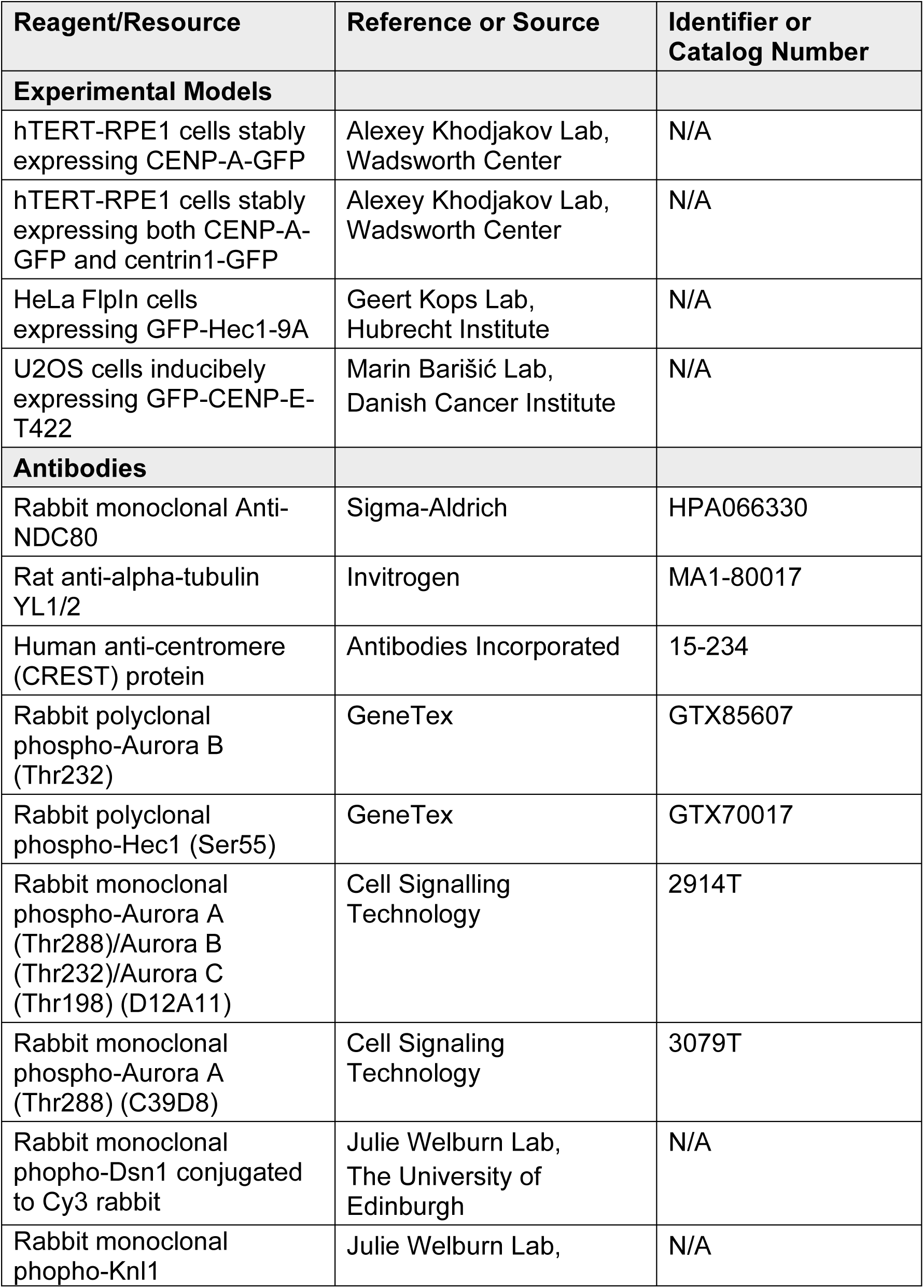
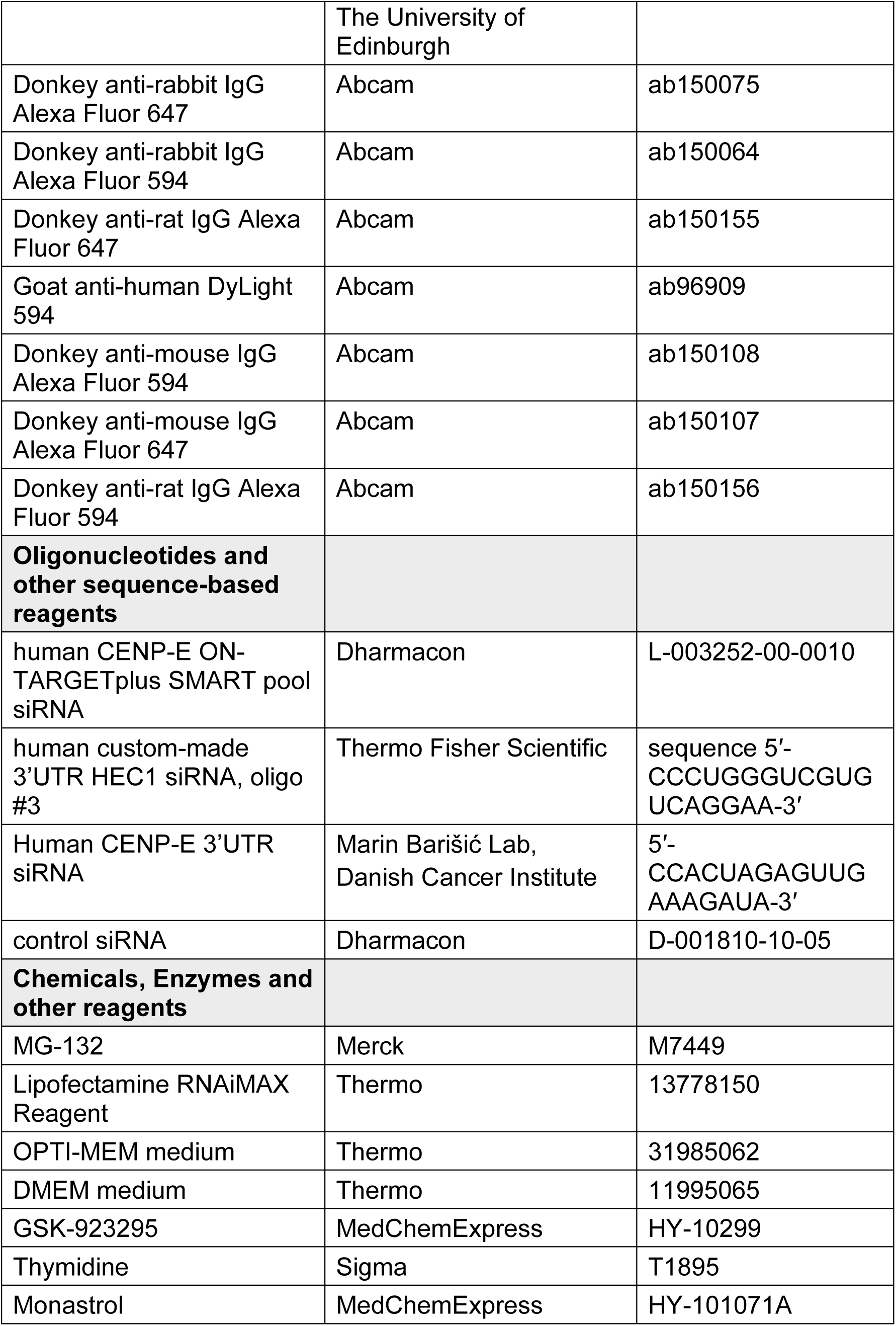
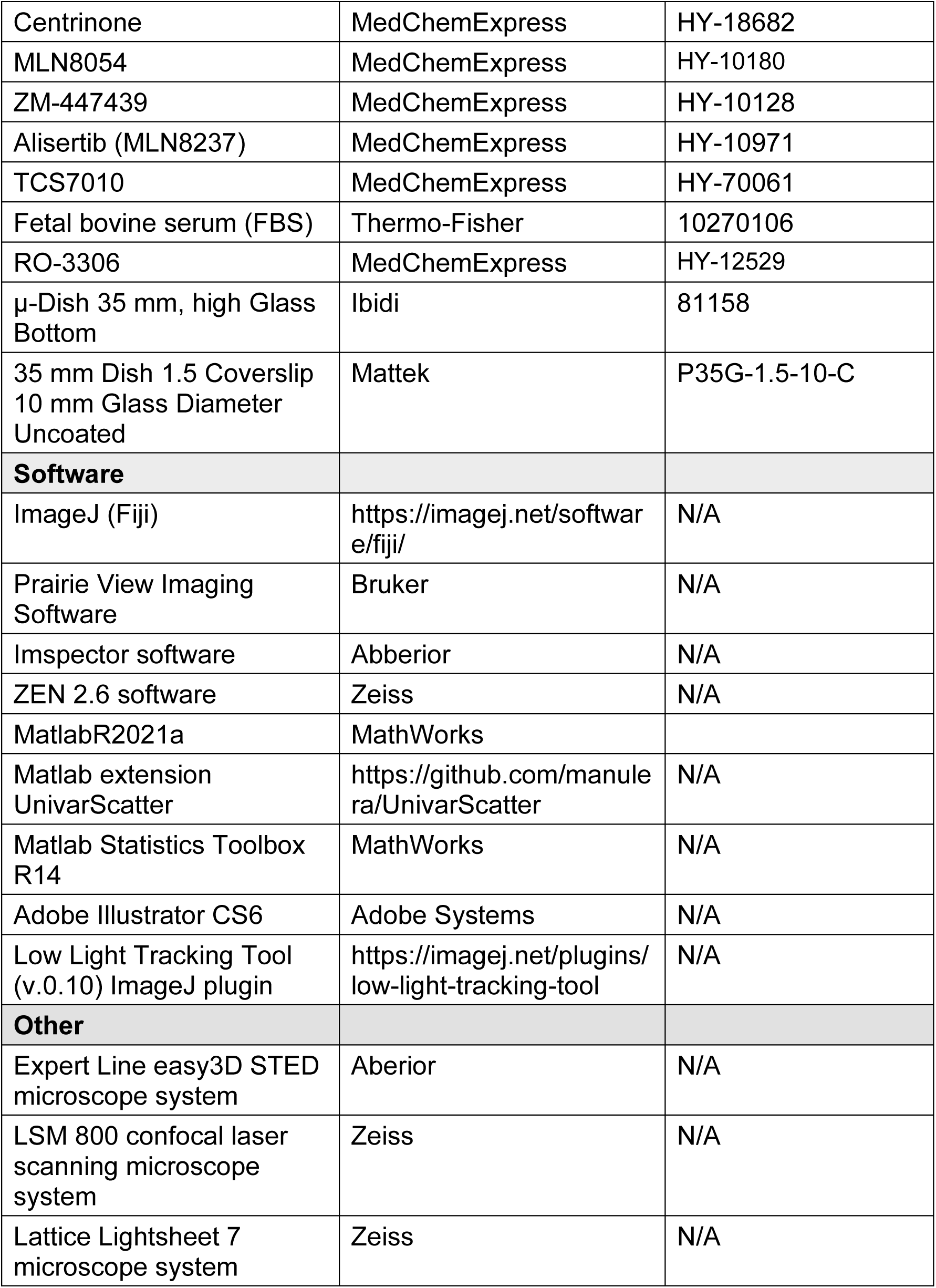

### Methods and Protocols

#### Cell lines and culture

Experiments were carried out using human hTERT-RPE1 (hTERT-immortalized retinal pigment epithelium) cells stably expressing CENP-A-GFP, human hTERT-RPE1 cells stably expressing both CENP-A-GFP and centrin1-GFP and human hTERT-RPE1 cells stably expressing CENP-A-GFP and Mis12-mCherry, all courtesy of Alexey Khodjakov (Wadsworth Center, New York State Department of Health, Albany, NY, USA), human HeLa (cervical carcinoma patient) cells expressing GFP-Hec1-9A, courtesy of Geert Kops (Hubrecht Institute, Utrecht, The Netherlands), and human U2OS (osteosarcoma patient) cells with inducible expression of phosphonull CENP-E mutated at the AurA/B-specific phospho-site Threonine 422 (T422A), which was a gift from Marin Barišić (Danish Cancer Institute, Copenhagen, Denmark). All cell lines were cultured in flasks in Dulbecco’s Modified Eagle’s Medium with 1 g/L D-glucose, pyruvate and L-glutamine (DMEM, Thermo Fisher, 11995065), supplemented with 10% (vol/vol) heat-inactivated Fetal Bovine Serum (FBS, Sigma-Aldrich, St. Louis, MO, USA) and penicillin (100 IU/mL)/streptomycin (100 mg/mL) solution (Lonza, Basel, Switzerland). The cells were kept at 37 °C and 5% CO2 in a humidified incubator (Galaxy 170 S CO2, Eppendorf, Hamburg, Germany) and regularly passaged at the confluence of 70-80%. None of the cell lines were authenticated. All cell lines have also been tested for mycoplasma contamination once a month by examining samples for extracellular DNA staining with SiR-DNA (100 nM, Spirochrome, Stein am Rhein, Switzerland) and Hoechst 33342 dye (1 drop/2 ml of NucBlue Live ReadyProbes Reagent, Thermo Fisher Scientific, Waltham, MA, USA) and have been confirmed to be mycoplasma free.

#### Sample preparation and siRNAs

At 80% confluence, the DMEM medium was removed from the flask and the cells were washed with 5 ml of phosphate buffered saline (PBS). Then, 1 ml 1% trypsin/ethylenediaminetetraacetic acid (EDTA, Biochrom AG, Berlin, Germany) was added to the flask and cells were incubated at 37 °C and 5% CO_2_ in a humidified incubator for 5 min. After incubation, trypsin was blocked by adding 4 ml of DMEM medium. For the RNAi experiments, cells were seeded to reach 60% confluence the next day and cultured on 35 mm uncoated plates with 0.17 mm (#1.5 coverglass) glass thickness (MatTek Corporation) in 1 ml of DMEM medium with the supplements described above. After one day of growth, cells were transfected with either targeting or non-targeting siRNA constructs which were diluted in OPTI-MEM medium (Thermo-Fisher) to a final concentration of 100 nM in the medium with cells. All transfections were performed 48 h before imaging using Lipofectamine RNAiMAX Reagent (Life Technologies, Waltham, MA, USA) according to the instructions provided by the manufacturer, unless otherwise indicated. Codepletions of Hec1 and CENP-E in HeLa cells were performed for two times in two subsequent days by the same protocol. After four hours of siRNA treatment, the medium was changed to the prewarmed DMEM medium. GFP-Hec1-9A construct in HeLa cells was expressed by addition of 1 μg ml^−1^ doxycycline for 24 h prior to fixation. Proteasome inhibitor MG-132 (Merck, M7449, IC50 value 100 nM) at a final concentration of 1 μM, was added 5 h before fixation for experiments involving HeLa GFP-Hec1-9A cells. Synchronization of GFP–Hec1-9A-expressing cells was achieved by treating cells seeded at 60% confluency with 2 mM thymidine (Sigma) for 24 hours, followed by four washes with pre-warmed DMEM. Cells were then incubated with 5 μM RO-3306 (MedChemExpress) for 16 hours and subsequently washed three times for 5 minutes each at 37 °C with pre-warmed DMEM. After release, cells were incubated for 2 hours in fresh medium, then treated with 1 μM MG-132 (Merck) for 2 hours prior to fixation.

The siRNA constructs used were: human CENP-E ON-TARGETplus SMART pool siRNA (L-003252-00-0010, human custom-made 3’UTR HEC1 siRNA (oligo #3, Thermo Fisher Scientific; sequence 5′-CCCUGGGUCGUGUCAGGAA-3′), and control siRNA (D-001810-10-05, Dharmacon, Lafayette, CO, USA). For experiments in U2OS cells expressing different CENP-E variants, endogenous CENP-E was depleted by transfecting cells with 20 nM 3′UTR-targeting siRNA (5′-CCACUAGAGUUGAAAGAUA-3′) 24 hours prior to fixation^23^. GFP-CENP-E expression was induced by adding doxycycline (1 μg/ml, Sigma-Aldrich) overnight. 1 μM MG-132 (Merck) was added for 2 hours in fresh medium before the fixation.

#### Drug treatments and drug washouts

CENP-E inhibitor GSK-923295 (MedChemExpress, Monmouth Junction, NJ, USA, IC50 value 3.2 nM) at a final concentration of 80 nM for RPE-1 and U2OS cells, was added 1-4 hours before imaging, or when noted acutely before imaging. Eg5 inhibitor Monastrol (HY-101071A/CS-6183, MedChemExpress, IC_50_ value 50 μM) at a final concentration of 100 μM, was added 3 hours before imaging or acutely before imaging when noted. Aurora A inhibitor MLN8054 (MedChemExpress, IC_50_ value 4 nM) at a final final concentration of 125 nM or 250 nM as noted, was added acutely before imaging or 30-60 minutes before fixation. Aurora A inhibitor Alisertib (MLN8237, MedChemExpress, IC_50_ value 1.2 nM) at a final final concentration of 125 nM, was added 30 minutes before fixation. Aurora A inhibitor TCS7010 (MedChemExpress, IC_50_ value 3.4 nM) at a final final concentration of 100 nM, was added 30 minutes before fixation. Aurora B inhibitor ZM-447439 (MedChemExpress, IC_50_ value 130 nM) at a final concentration of 3 μM, was added 15 minutes before fixation. PLK4 inhibitor Centrinone (MedChemExpress, K_i_ value 0.16 nM) at a final concentration of 300 nM, was added 1-5 days before the imaging as noted in the manuscript. The stock solutions for all drugs were prepared in DMSO. The stock solutions of all drugs were kept aliquoted at 10-50 μL at −20 °C for a maximum period of three months or at −80 °C for a maximum period of six months. New aliquots were thawed weekly for each new experiment. All drugs were added to DMEM media used for cell culture to obtain the final concentration of a drug as described. Drug washouts were performed by replacing drug-containing medium with 2 mL of fresh pre-warmed DMEM medium followed by four subsequent washouts with 2 mL of pre-warmed DMEM.

To deplete centrioles, Plk4 kinase activity was inhibited with centrinone, a small molecule inhibitor that blocks the centriole duplication cycle, and varied the duration of treatment to obtain spindles with different numbers of centrioles on each centrosome, a method established previously^19^. Subsequently, we treated Plk4-inhibited cells with CENP-E inhibitor or DMSO and imaged cells a few hours after treatment using the same live cell imaging protocol described above. After acute treatment of CENP-E inhibited cells with the Eg5 inhibitor monastrol (50 uM), spindles quickly shortened, as expected from previous reports^70^. Similarly, spindle shortening was observed in most CENP-E inhibited cells after acute addition of the Aurora A inhibitor MLN8237 (125 nM), as expected from previous studies^50^.

Regarding cells imaged by confocal microscopy after the chronic inhibition of CENP-E by GSK-923295 (80 nM), all pseudo-metaphase spindles chosen for imaging were phenotypically similar between each other and between conditions where CENP-E was depleted or reactivated after washout of the CENP-E inhibitor: 1) in all cells few chromosomes were localized close to one of the spindle poles, called polar chromosomes, and their numbers ranged from 2-16 per cell; 2) occasionally in some cells few chromosomes were localized already in between spindle pole and equator; and 3) in all cells most chromosomes were already aligned and tightly packed at the metaphase plate. This type of chromosome arrangement was expected from previously published data that reported that only 10-30% of chromosomes upon NEBD require CENP-E-mediated alignment^8^. Likewise, under all conditions where CENP-E was perturbed and cells imaged by confocal microscopy during pseudo-metaphase, the overall displacement of spindle poles from each other was negligible (data not shown), consistent with cells being in a pseudo-metaphase state where there is no net change in spindle length. For LLSM-based assay, no inclusion criteria for imaging were used as the cells were non-synchronized and entered mitosis stochastically. Only randomly selected cells that entered mitosis and subsequently entered anaphase during imaging time were used for kinetochore and centrosome tracking analysis.

#### Immunofluorescence

For cell labeling with phopho-Knl1, phopho-Dsn1, phospho-Hec1, phospho-Aurora A, phospho-Aurora B, phospho-AuroraA/B/C, RPE-1 cells expressing CENP-A-GFP and centrin1-GFP grown on glass-bottom dishes (14 mm, No. 1.5, MatTek Corporation, Darmstadt, Germany) were pre-extracted with pre-warmed PEM buffer at 37 °C (0.1 M PIPES, 0.001 M MgCl2 × 6 H2O, 0.001 M EDTA, 0.5% Triton-X-100) for 30 s at room temperature and fixed by 1 ml of pre-warmed solution of 4% paraformaldehyde (PFA) for 20 min. All other antibodies were used on cells fixed for 2 minutes with ice-cold methanol, except for those used in STED microscopy, which are described below. After fixation, cells were washed 3 times for 5 min with 1 ml of PBS and permeabilized with 0.5% Triton-X-100 solution in water for 30 min at room temperature. To block unspecific binding, cells were incubated in 1 ml of blocking buffer (2% bovine serum albumin, BSA) for 2 h at room temperature. The cells were then washed 3 times for 5 min with 1 ml of PBS and incubated with 500 µl of primary antibody solution overnight at 4°C. Antibody incubation was performed using a blocking solution composed of 0.1% Triton, 1% BSA in PBS.

Following primary antibodies were used: rabbit monoclonal phopho-Knl1 (1.5 mg/ml, gift from J. Welburn, diluted 1:1000), rabbit monoclonal phopho-Dsn1 conjugated to Cy3 (0.5 mg/ml, gift from J. Welburn, diluted 1:250, diluted 1:500), rabbit monoclonal phospho-Aurora A (Thr288) (C39D8) (3079T, Cell Signaling Technology, diluted 1:500), rabbit monoclonal phospho-Aurora A (Thr288)/Aurora B (Thr232)/Aurora C (Thr198) (D12A11) (2914T, Cell Signalling Technology, diluted 1:500), rabbit polyclonal phospho-Hec1 (Ser55) (GTX70017, GeneTex, diluted 1:500), rabbit polyclonal phospho-Aurora B (Thr232) (GTX85607, GeneTex, diluted 1:500), rabbit monoclonal Anti-NDC80 (HPA066330-100uL, Sigma-Aldrich, diluted 1:250), human anti-centromere (CREST) protein (15-234, Antibodies Incorporated, 1:500), and rat anti-alpha-tubulin YL1/2 (MA1-80017, Invitrogen, diluted 1:500). Where indicated, DAPI (1 µg/mL) was used for chromosome visualization.

After primary antibody, cells were washed in PBS and then incubated in 500 μL of secondary antibody solution for 1 h at room temperature. Following secondary antibodies were used: Donkey anti-rabbit IgG Alexa Fluor 647 (ab150075, Abcam, diluted 1:1000) for phopho-Dsn1, Donkey anti-rabbit IgG Alexa Fluor 594 (ab150064, Abcam, diluted 1:1000) for all other rabbit antibodies, Donkey anti-mouse IgG Alexa Fluor 594 (ab150108, Abcam, diluted, 1:1000) for all mouse antibodies, goat anti-human DyLight 594 (Abcam, ab96909, diluted 1:1000), and Donkey anti-rat IgG Alexa Fluor 647 (ab150155, Abcam, diluted 1:500). Finally, cells were washed with 1 ml of PBS, 3 times for 10 min. Cells were imaged either immediately following the imaging or were kept at 4°C before imaging for a maximum period of one week.

To visualize alpha-tubulin in super-resolution by STED and Airyscan microscopy in the RPE-1 CENP-A-GFP centrin1-GFP cell line, the ice-cold methanol protocol was avoided because it destroyed the unstable fraction of microtubules^71^. Cells were washed with cell extraction buffer (CEB) and fixed with 3.2% paraformaldehyde (PFA) and 0.1% glutaraldehyde (GA) in PEM buffer (0.1 M PIPES, 0.001 M MgCl2 × 6 H2O, 0.001 M EGTA, 0.5% Triton-X-100) for 10 min at room temperature. After fixation with PFA and GA, for quenching, cells were incubated in 1 ml of freshly prepared 0.1% borohydride in PBS for 7 min and then in 1 mL of 100 mM NH_4_Cl and 100 mM glycine in PBS for 10 min at room temperature. To block nonspecific binding of antibodies, cells were incubated in 500 μL blocking/permeabilization buffer (2% normal goat serum and 0.5% Triton-X-100 in water) for 2 h at room temperature. Cells were then incubated in 500 μL of primary antibody solution containing rat anti-alpha-tubulin YL1/2 (MA1-80017, Invitrogen, diluted 1:300) overnight at 4 °C. After incubation with a primary antibody, cells were washed 3 times for 10 min with 1 ml of PBS and then incubated with 500 µl of secondary antibody containing donkey anti-rat IgG Alexa Fluor 594 (ab150156, Abcam, diluted 1:300) for 2 h at room temperature, followed by wash with 1 ml of PBS 3 times for 10 min.

#### Imaging

The STED confocal microscope system (Abberior Instruments) and the LSM800 laser scanning confocal system (Zeiss) were used for the live-cell experiments we termed the confocal-based imaging assay. STED microscopy was also used to image all fixed cells in super-resolution. STED microscopy was performed using an Expert Line easy3D STED microscope system (Abberior Instruments, Göttingen, Germany) with the 100x/1.4NA UPLSAPO100x oil objective (Olympus, Tokio, Japan) and an avalanche photodiode detector (APD). The 488 nm line was used for excitation, with the addition of the 561 nm line for excitation and the 775 nm laser line for depletion during STED super-resolution imaging. The images were acquired using Imspector software. The xy pixel size for fixed cells was 20 nm and 10 focal planes were acquired with a 300 nm distance between the planes. For confocal live cell imaging of cells, the xy pixel size was 80 nm and 16 focal images were acquired, with 0.5 µm distance between the planes and 1min time intervals between different frames. During imaging, live cells were kept at 37 °C and at 5% CO_2_ in the Okolab stage incubation chamber system (Okolab, Pozzuoli, NA, Italy). Live-cell imaging following acute reactivation or depletion of CENP-E (Extended Data Fig. 1) was performed using an Opterra I point-scanning confocal system (Bruker), as previously described^11^.

LSM 800 confocal laser scanning microscope system (Zeiss) was used for the rest of fixed-cell confocal-based imaging in hTERT-RPE1 cells expressing CENP-A-GFP and centrin1-GFP with the following parameters: sampling in xy, 0.27 µm; z step size, 0.5 µm; total number of slices, 32; pinhole, 48.9 µm; unidirectional scan speed, 10; averaging, 2; 63x Oil DIC f/ELYRA objective (1.4 NA), 488 nm laser line (0.1-1% power for different experiments), and detection ranges of 450–558 nm for the green channel, 561 nm laser line (0.1-1% power for different experiments) and detection range of 565-650 nm for the red channel, 640 nm laser line (0.1-1% power for different experiments), and detection range of 656-700 nm for the far red channel, and 405 nm laser line (0.5% power) and detection range of 400-450 for the blue channel. Images were acquired using ZEN 2.6 (blue edition; Zeiss). Imaging was done on multiples positions determined by the user simultaneously, and at the confocal lateral resolution. For imaging of tubulin in super-resolution following acute Aurora A inhibition in CENP-E-inhibited cells, Airyscan mode of LSM 800 was used. Following parameters were used: sampling in xy, 2 µm; z step size, 0.146 µm; total number of slices, 41; pinhole, 265 µm; unidirectional scan speed, 7; averaging, 2; 63x Oil DIC f/ELYRA objective (1.4 NA), 488 nm laser line (1.5% power), 561 nm laser line (1% power) and detection ranges of 450–580 and 565-700, for the green and red channels, respectively. Images were acquired using ZEN 2.6 (blue edition; Zeiss). Imaging was done on multiples positions determined by the user.

The Lattice Lightsheet 7 microscope system (Zeiss) was used for live cell imaging of hTERT-RPE1 cells expressing CENP-A-GFP and Centrin1-GFP in assay we termed the LLSM-based imaging assay. The system was equipped with an illumination objective lens 13.3×/0.4 (at a 30° angle to cover the glass) with a static phase element and a detection objective lens 44.83×/1.0 (at a 60° angle to cover the glass) with an Alvarez manipulator. Images were acquired using ZEN 2.6 (blue edition; Zeiss). The automatic immersion of water was applied from the motorized dispenser at an interval of 20 or 30 minutes. Right after sample mounting, four steps of the ‘create immersion’ auto immersion option were applied. The sample was illuminated with a 488-nm diode laser (power output 10 Mw) with laser power set to 1-2%. The detection module consisted of a Hamamatsu ORCA-Fusion sCMOS camera with exposure time set to 15-20 ms. The LBF 405/488/561/642 emission filter was used. During imaging, cells were kept at 37 °C and at 5% CO2 in a Zeiss stage incubation chamber system (Zeiss). The width of the imaging area in the x dimension was set from 1 to 1.5 mm.

#### Tracking of centrosomes and kinetochores

The spatial x and y coordinates of the kinetochore pairs were extracted in each time frame using the Low Light Tracking Tool (v.0.10), an ImageJ plugin based on the Nested Maximum Likelihood Algorithm, as previously described^70^. Tracking of polar kinetochores in the x and y planes was performed on the maximum intensity projections of all acquired z planes. Some kinetochore pairs could not be successfully tracked in all frames, mainly owing to cell and spindle movements in the z-direction over time. Spindle poles were manually tracked with points placed between the center of the two centrioles or centriole in centrinone treated cells. Kinetochore pairs that were aligned at the start of the imaging in a confocal-based imaging assay were also manually tracked in two dimensions. Quantitative analysis of all parameters was performed using custom-made MATLAB scripts (MatlabR2021a 9.10.0) scripts. Due to the inability to reliably track polar kinetochores across all time frames due to neighboring kinetochores, tracking of polar kinetochores routinely commenced a few frames before the kinetochore pair began moving towards the equatorial plane. In the CENP-E reactivated and CENP-E-depleted RPE-1 cells imaged by confocal microscopy, kinetochores were tracked from the onset of imaging until the successful identification of the same kinetochore pair, which occurred when the kinetochore pair was lost in the imaging plane, among other kinetochores, or when the imaging session ended, whichever came first.

#### Quantification of mean pHec1, pKnl1 and pAuroraB intensities on kinetochore pairs and pAuroraA on spindle poles

The fluorescence intensity signal of each kinetochore in both CENP-A and respective kinetochore protein channel (pDsn1, pKnl1) was measured by using the “*Oval selection*” tool in ImageJ with the size and position defined by the borders of the CENP-A signal of each kinetochore in the sum intensity projection of all acquired z-planes. pAurora A signal was measured using the “*Oval selection*” tool around the centriole signal as the center or around the signal of the Aurora A of the same size on the pole without centriole. The background fluorescence intensity measured in the cytoplasm was subtracted from the obtained values, and the calculated integrated intensity value was divided by the number of z-stacks used to generate the sum projection of each cell. The obtained mean intensity value subtracted for background for each kinetochore was normalized to the intensity of the CENP-A signal. For the pHec1 labeling experiments, the mean signal of Hec1 and CENP-A was measured by the same approach but on one z-plane where kinetochore CENP-A signal was the highest. The mean signal of the pAurora B was measured between extent of CENP-A signals of sister kinetochore pairs, and the mean value was normalized to mean value of CENP-A signal of each sister kinetochore pair. All movies used for the analysis were performed using the same fixation protocol and were imaged with the same imaging parameters in at least three independent biological replicates. To avoid measurement of non-specific labeling of the spindle poles that are known artefacts of phospho-antibodies to kinetochore proteins (pAurB, pKnl1, pHec1)^30^, only the kinetochores outside of the circular pole signal (∼1 μm from the centrosome) were measured and used for analysis.

#### Quantification of mean signal intensities of proteins after RNAi interreference

The fluorescence intensity signals were measured in triplicate samples for targeting and non-targeting groups for each siRNA treatment. Each siRNA targeting and non-targeting triplicates were imaged by the same protocol. Proteins were measured in mitotic cells at their respective locations during mitosis by using the “*Polygon selection tool*”, as established by many previous studies: CENP-E and Hec1 were measured in area occupied by chromosomes, as defined by the DAPI on sum of all acquired planes.

#### Quantification of polar kinetochore pairs

In all experiments, polar kinetochores were defined as kinetochore pairs positioned closer to one of the spindle poles than to the center of the equatorial plane, with all others classified as aligned. In fixed-cell images, the polar category included both kinetochores near the pole and those in transit toward the metaphase plate. Kinetochore pairs with at least one kinetochore located 3 µm or less from the equatorial plane were assigned to the aligned group. In confocal-based experiments polar chromosomes were defined from the onset of imaging. In cells treated with centrinone and their respective DMSO controls, the number of polar chromosomes was measured in time when the spindle finished the bipolarization process, e.g., prometaphase elongation of the spindle. This was done because all spindles of the 1:0 type with only one centriole at the beginning of mitosis went into the monopolar state and stayed in this state for variable durations before bipolarization, ranging from 10 to 50 minutes, as reported previously^26^. The residence time of polar chromosomes on spindle poles, i.e., the efficiency of congression for each pole, was defined as the time from spindle bipolarization until the polar chromosome either entered the metaphase plate or the cell entered anaphase with an uncongressed polar chromosome. Duration of mitosis was defined as the time from the first signs of nuclear envelope breakdown to one frame before visible onset of anaphase defined by the separation of sister kinetochore pairs. In fixed cells, the number of polar chromosomes per single cell, as defined above, was used as a readout of congression efficiency.

Congression velocity was calculated for each pair of kinetochores in the last 6 minutes before the center of the pair of kinetochores surpassed 2 µm from the equatorial plane measured as the nearest distance from a center of a pair to a plane. These times represent the fast phase of kinetochore movement towards the plate^11^. Aligned kinetochore pair was defined as every pair that was found 3 μm from the equatorial plane at any given time. The equatorial or metaphase plane was defined in each time frame as the line perpendicular to the line connecting the centrosomes at their midpoint. The angle between the kinetochore pairs and the main spindle axis was defined as an angle between a line connecting two centrosomes and a line connecting the center of the kinetochore pair and a spindle pole nearest to the respective pair. The angle between the midpoint of sister kinetochores and the centrosome-to-centrosome axis was measured using the angle tool in ImageJ. The successful alignment of a polar kinetochore pair to the metaphase plate was defined as the moment when the midpoint between sister kinetochores first crossed within 2 μm of the equatorial plane.

Maximum metaphase plate spread was measured in ImageJ as the distance between the outermost chromosome boundaries along the main spindle axis, based on the DAPI signal. The number of polar chromosomes was determined by counting kinetochore pairs in CREST-labeled cells or individual chromosomes in DAPI-labeled cells. In Hec1-9A overexpression experiments, no selection bias was applied for Hec1-9A–expressing cells; however, the proportion of expressing cells was consistent across experiments, accounting for approximately 10% of imaged cells. Similarly, in CENP-E-T422 overexpression experiments, no selection bias was applied, and CENP-E-T422–expressing cells represented less than ∼5% of imaged cells. Multipolar spindles, identified by tubulin or centrin staining, were excluded from analysis. Spindle length was measured in ImageJ as the distance between centriole pairs in 2D maximum projections. The distance to the nearest centrosome or to the equatorial plane was measured from the midpoint between sister kinetochores to either the center of the nearest centriole pair or the equatorial plane, respectively. Interkinetochore distance was measured as the distance between the centers of two sister kinetochores.

#### Image processing and statistical analysis

Image processing was performed in ImageJ (National Institutes of Health, Bethesda, MD, USA). Quantification and statistical analysis were performed in MatLab. The figures were assembled in Adobe Illustrator CS6 (Adobe Systems, Mountain View, CA, USA). Raw images were used for quantification. The images of spindles were rotated in every frame to fit the long axis of the spindle to be parallel with the central long axis of the box in ImageJ and the spindle short axis to be parallel with the central short axis of the designated box in ImageJ. The designated box sizes were cut to the same dimensions for all panels in the figures where the same experimental setups were used across the treatments. When comparing different treatments in channels in which the same protein was labeled, the minimum and maximum intensity of that channel was set to the values in the control treatment. When indicated, the smoothing of the images was performed using the “Gaussian blur” function in ImageJ (s=0.5-1.0). Colour-coded maximum intensity projections of the z-stacks were done using the “*Temporal colour code*” tool in Fiji by applying “16-color” or “Spectrum” lookup-table (LUT) or other LUT as indicated. For the generation of univariate scatter plots, the open Matlab extension “UnivarScatter” was used.

Data are given as mean ± standard deviation (s.t.d.), unless otherwise stated. The mean line was plotted to encompass a minimum of 60% of the data points for each treatment. Other dispersion measures used are defined in their respective figure captions or in Fig. 1 and Extended Data Fig. 1 if the same measures are used across all figures. The exact values of n are given in the respective figure captions, where n represents the number of cells or the number of tracked kinetochore pairs, as defined for each n in the figure captions or tables. The number of independent biological replicates is also given in the figure captions. An independent experiment for acute drug treatments was defined by the separate addition of a drug to a population of cells in a dish. The number of cells imaged simultaneously ranged from 1 to 7, depending on the specific microscopy system used. The p values when comparing data from multiple classes that followed a normal distribution were obtained using the one-way ANOVA test followed by Two-sided Tukey’s Honest Significant Difference (HSD) test (significance level was 5%). p < 0.05 was considered statistically significant, very significant if 0.001 < p < 0.01 and extremely significant if p < 0.001. Values of all significant differences are given with the degree of significance indicated (∗0.01 <p < 0.05, ∗∗0.001 < p < 0.01, ∗∗∗p < 0.001, ∗∗∗∗ < 0.0001). For the linear regression correlation measure between two parameters, the nonparametric Spearman correlation coefficient, termed rs, was used where p<0.001, calculated using the ‘corr’ function in Matlab (Statistics Toolbox R14). No statistical methods were used to predetermine the sample size. The experiments were not randomized and, except where stated, the investigators were not blinded to allocation during experiments and outcome evaluation.

## DATA AVAILABILITY

Source data for the main figures is provided with this paper. All other data supporting the findings of this study are available from the corresponding authors on reasonable request.

## CODE AVAILABILITY

Codes used to analyze and plot the data are available from the corresponding authors on request.

## EXTENDED DATA FIGURE LEGENDS

**Extended Data Fig. 1.**
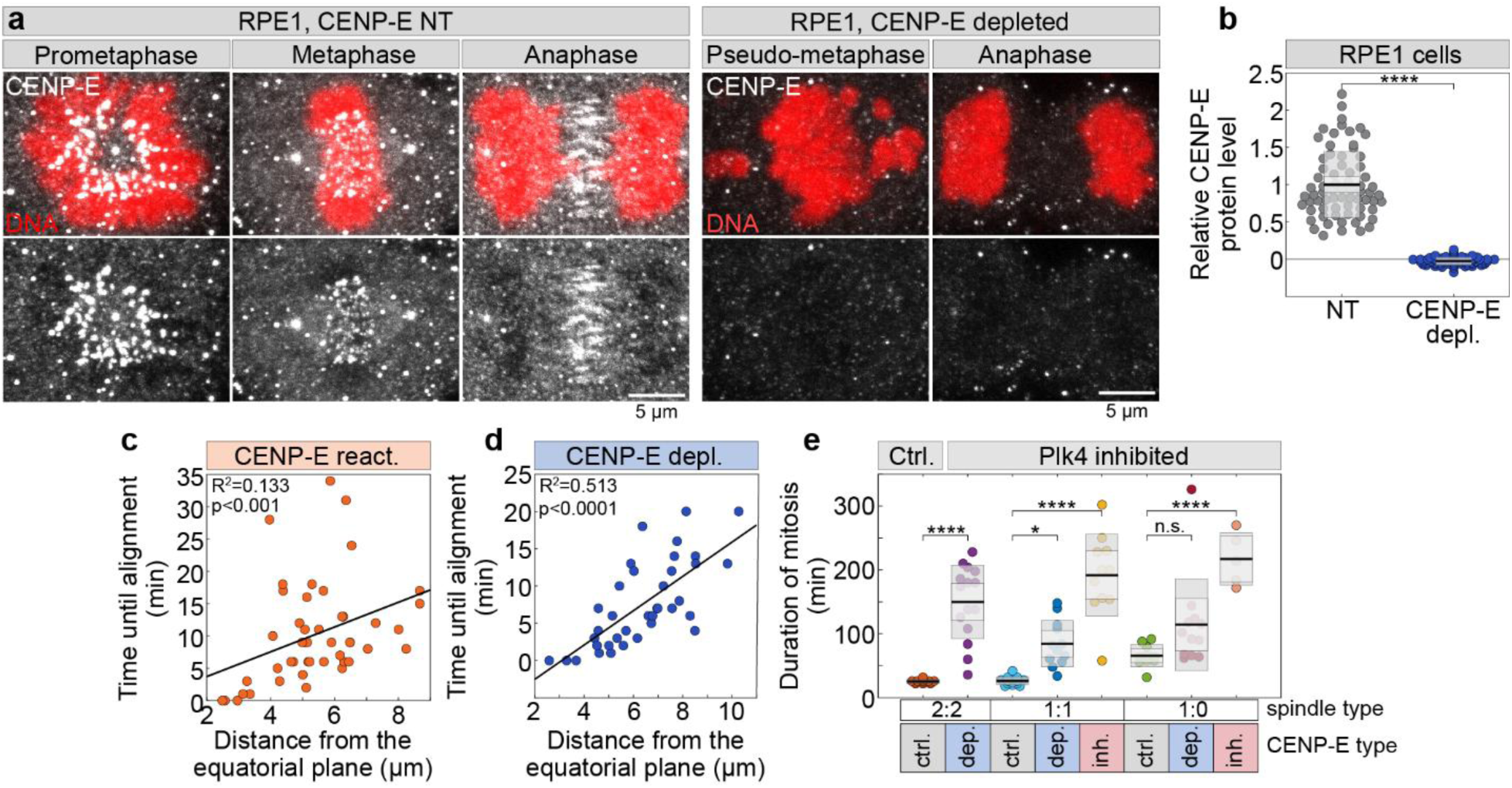
Speed of congression initiation is correlated with the distance of the polar chromosome from the centrosome. (**a**) Representative images of RPE-1 cells treated with non-targeting (NT) siRNAs (left) or CENP-E siRNA (right), immunostained with antibodies against CENP-E at different mitotic stages as indicated. Cells were wild-type (WT) and stained with DAPI (red). Top panels show maximum intensity projections with merged channels; bottom panels show the CENP-E signal (gray). (**b**) Quantification of CENP-E protein levels after its depletion normalized to the non-targeting control group. Numbers: 62 and 57 cells, respectively, each from three independent experiments. (**c**, **d**) Correlation between the initial distance of polar kinetochore pairs to the equatorial plane and their total congression time, following washout of CENP-E inhibitor (c) or CENP-E depletion (d); lines represent regression fits. Numbers: reactivated (18 cells, 86 kinetochore pairs), CENP-E depleted (18 cells, 69 kinetochore pairs), each from more than 3 independent experiments. (**e**) Duration of mitosis, time from nuclear envelope breakdown to anaphase onset, in cells with different numbers of centrioles and under different treatments, as indicated. Black lines indicate means; light and dark gray areas represent 95% confidence intervals and standard deviations, respectively. Numbers for e given in Fig. 1. Statistics: ANOVA with post-hoc Tukey’s HSD test. Symbols indicate: n.s., P > 0.05; **, P ≤ 0.01; ***, P ≤ 0.001; ****, P ≤ 0.0001.

**Extended Data Fig. 2.**
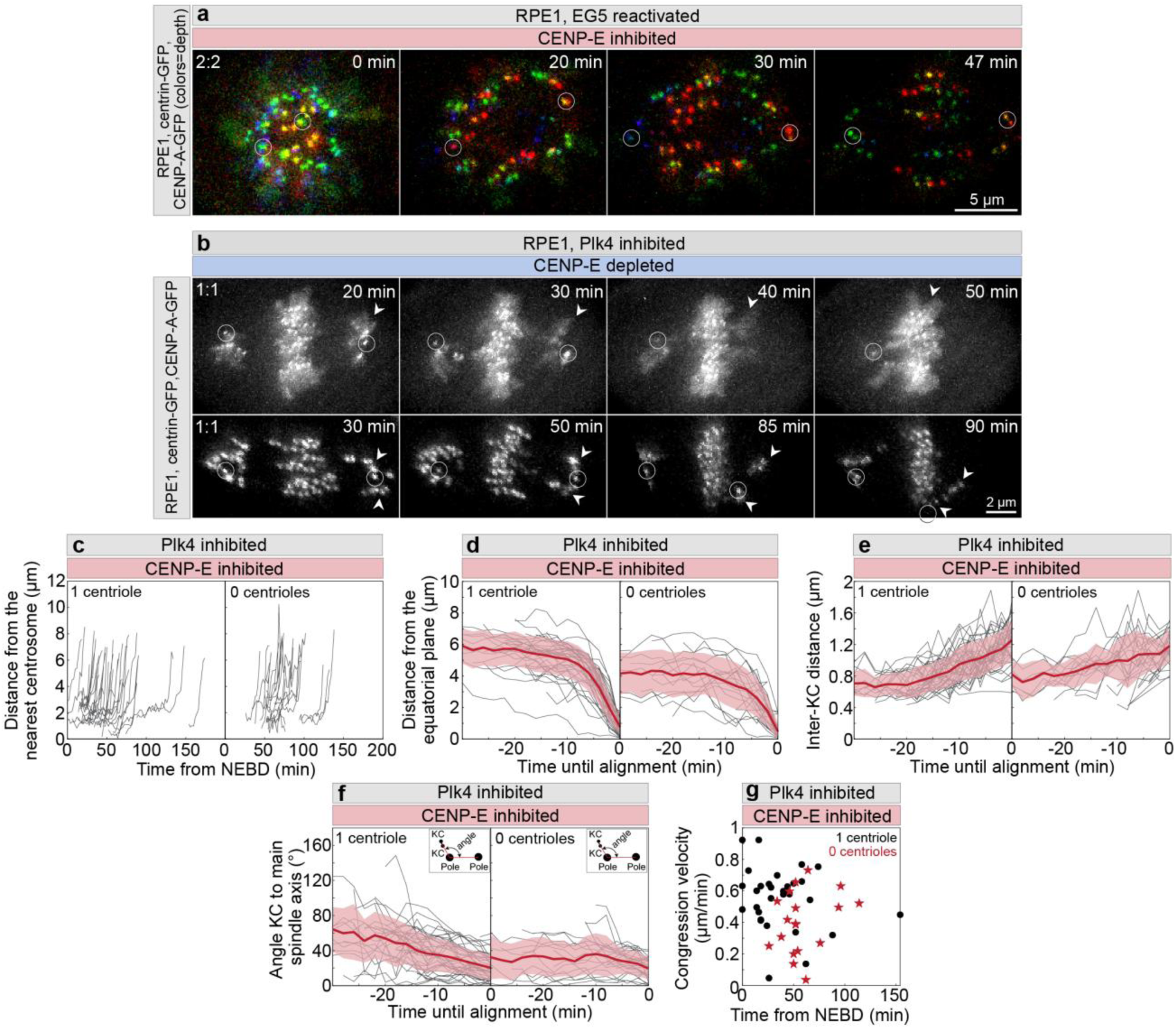
Chromosome movement during congression is similar in cells with different numbers of centrioles. (**a**) Representative images of an RPE1 cell expressing CENP-A-GFP and centrin1-GFP, showing different time points following the washout of the Eg5 inhibitor monastrol and addition of 80 nM CENP-E inhibitor GSK923295. Time 0 marks the point of monastrol washout and GSK923295 addition. Centrioles are circled. (**b**) Representative time frames of two RPE1 cells expressing CENP-A-GFP and centrin1-GFP following CENP-E depletion, starting at NEBD. Centrioles are circled in white; arrowheads indicate polar chromosomes that align after centriole separation from the centrosome. (**c**–**f**) Quantification of kinetochore behavior from nuclear envelope breakdown (NEBD) until successful alignment for initially polar kinetochore pairs in the presence of CENP-E inhibitor: distance from the nearest spindle pole (c), distance to the equatorial plane (d), interkinetochore (inter-KC) distance (e), and angle relative to the main spindle axis (f, schematic shown in graph). Data compare chromosomes congressing from spindle poles with one centriole (left) and without centrioles (right). Thick lines indicate mean values; shaded areas represent standard deviations. (**g**) Velocity of chromosome congression during the 6-minute window preceding full alignment, plotted against the time from NEBD when the congression event began, for polar chromosomes from poles with one centriole (black) or without centrioles (red). Numbers of cells and independent experiments for (c-g) given in Fig. 1. Symbols indicate: KC, kinetochore; NEBD, nuclear envelope breakdown.

**Extended Data Fig. 3.**
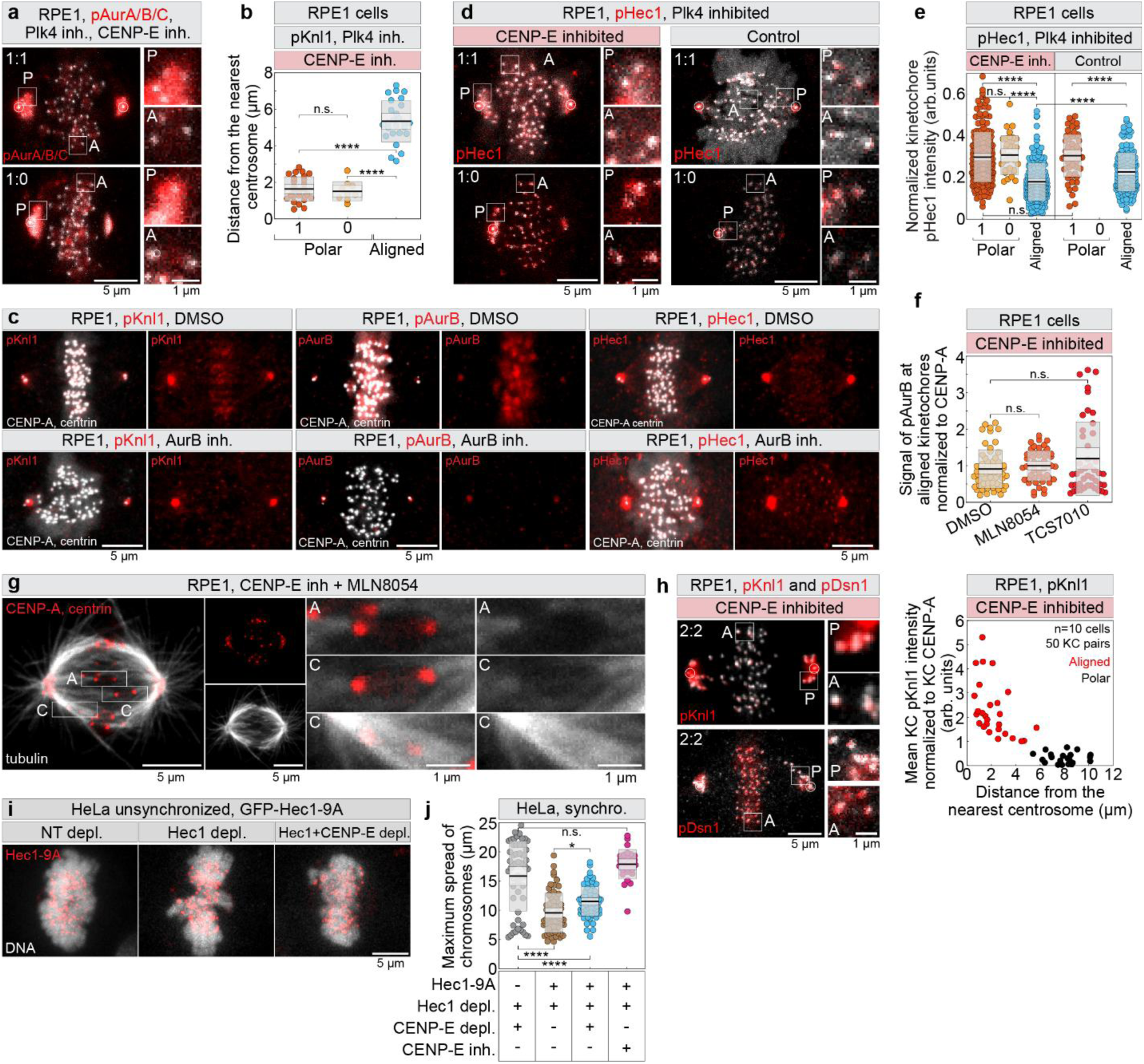
Aurora B-mediated phosphorylation gradually declines on outer kinetochore substrates during congression of polar chromosomes. (**a**) Representative images and enlarged insets of RPE-1 cells expressing CENP-A-GFP and centrin1-GFP (centrioles circled), immunostained for pT288-Aurora A, pT232-Aurora B, and pT198-Aurora C, showing spindles with different centriole numbers following continuous 300 nM centrinone and acute CENP-E inhibitor treatment as indicated. Images are maximum intensity projections. A, aligned; P, polar. (**b**) Distance from the nearest spindle pole for kinetochore groups identified in the pKnl1 immunolabelling experiment after CENP-E inhibition. Numbers: 62 kinetochore pairs, from two independent experiments. (**c**) Representative images of cells stained for phosphorylated kinetochore proteins (pS60-Knl1, pT232-Aurora B, pS55-Hec1, red), with separated red channels shown at right, following 15-minute treatment with DMSO or 3 μM Aurora B inhibitor ZM-447439. Images are maximum projections. (**d**) Representative images of cells expressing CENP-A-GFP and centrin1-GFP (grey), with centrioles circled in white, immunostained for phosphorylated Hec1 (pS55-Hec1, red) under the indicated conditions, with enlarged insets at right; images are maximum projections (**e**) Mean pS55-Hec1 intensity at kinetochores normalized to CENP-A intensity across treatments and centriole numbers. Numbers: 594 kinetochores, 67 cells, from more than 3 independent experiments. (**f**) Levels of pT232-Aurora B on aligned kinetochores normalized to average CENP-A levels in CENP-E–inhibited cells after indicated acute treatments. Numbers given on Fig. 4. (**g**) Representative image of an RPE-1 cell expressing CENP-A-GFP and centrin1-GFP (red), treated as indicated, showing congressing (C) and aligned (A) kinetochores with merged and separated channels, acquired by super-resolution Airyscan microscopy. Image is a deconvolved maximum projection of 5 z-planes, with α-tubulin stained in grey. (**h**) Representative images and enlarged insets of RPE-1 cells expressing CENP-A-GFP and centrin1-GFP, immunostained for pKnl1 (top left) and pDsn1 (pS100, pS109-Dsn1, bottom left) after CENP-E inhibition, and quantification of mean pKnl1 signal intensity versus spindle pole distance for polar and aligned kinetochore pairs (right). Experiment done in two independent experiments. (**i**) Representative examples of unsynchronized HeLa cells expressing GFP-Hec1-9A (red) with DNA stained by DAPI (grey) in treatments indicated on the top. (**j**) Maximum chromosome spread in synchronized HeLa cells under the indicated treatments, as shown in the legend. Numbers given of Fig. 5. Statistics: ANOVA with post-hoc Tukey’s HSD test. Symbols indicate: n.s., P > 0.05; ****, P ≤ 0.0001; inh., inhibited; KC, kinetochore.

## VIDEO LEGENDS

**Video 1. RPE-1 cells stably expressing CENP-A-GFP and Centrin1-GFP in continuously Plk4-inhibited cells with different numbers of centrioles following CENP-E inhibition or depletion.** RPE-1 cells expressing CENP-A-GFP and Centrin1-GFP were treated with a Plk4 inhibitor for 2 days to generate centrosomes with varying centriole numbers (1:1, 1:0, and 0:0). Cells were then subjected either to CENP-E inhibition with drug treatment for 3 hours prior to imaging (CENP-E inhibited, left) or to CENP-E depletion via siRNA for 48 hours (CENP-E depleted, right). Cells were imaged 3 hours after drug addition or 48 hours after siRNA transfection. Kinetochores and centrioles are depth color-coded using the 16-color LUT from blue (bottom) to red (top).

